# Resolving atomic-level dynamics and interactions of high molecular weight hyaluronic acid by multidimensional solid-state NMR

**DOI:** 10.1101/2024.02.29.582163

**Authors:** Pushpa Rampratap, Alessia Lasorsa, Abinaya Arunachalam, Marleen Kamperman, Marthe T.C. Walvoort, Patrick C.A. van der Wel

## Abstract

High molecular weight (HMW) hyaluronic acid (HA) is a highly abundant natural polysaccharide and a fundamental component of the extracellular matrix (ECM). Its size and concentration regulate tissues’ macro- and microenvironments, and its upregulation is a hallmark feature of certain tumors. Yet, the conformational dynamics of HMW-HA and how it engages with components of the ECM microenvironment remain poorly understood on the molecular level. Probing the molecular structure and dynamics of HMW polysaccharides in a hydrated, physiological-like environment is crucial but also technically challenging. Here, we deploy advanced magic-angle-spinning (MAS) solid-state NMR (ssNMR) spectroscopy in combination with isotopic enrichment to enable an in-depth study of HMW-HA to address this challenge. This approach resolves multiple coexisting HA conformations and dynamics as a function of environmental conditions. By combining ^13^C-labeled HA with unlabeled ECM components, we detect by MAS NMR HA-specific changes in global and local conformational dynamics as a consequence of hydration and ECM interactions. These measurements reveal atom-specific variations in dynamics and structure of the N-acetylglucosamine (GlcNAc) moiety of HA. We discuss possible implications for interactions that stabilize the structure of HMW-HA and facilitate its recognition by HA-binding proteins. The described methods apply similarly to studies of the molecular structure and dynamics of HA in tumor contexts and in other biological tissues, as well as HMW-HA hydrogels and nanoparticles used for biomedical and/or pharmaceutical applications.

## 1. Introduction

The extracellular matrix (ECM) is a complex network of macromolecules that envelops cells in tissues and organs. It serves as both a structural support and a contributor to tissue function ^1^. The ECM comprises various major components, including collagen, proteoglycans, glycoproteins, fibronectin, laminin, and hyaluronic acid (HA) polysaccharides (as illustrated in Figure 1A). It is now realized that the composition and mechanical properties of the ECM impact the development and behavior of nearby cells ^2^. A well-known example of the dramatic role of the ECM is in the context of cancer, where tumor tissues are characterized by a clear increase in ECM-related stiffness^3^. Notably, the biomechanical properties of the ECM are seen as important factors in cancer development and progression. Thus, there is much interest in studying the molecular underpinnings of the (biomechanical) role of the ECM and its constituents. Although much emphasis has been placed on studies of proteinaceous components, polysaccharides like HA are clearly crucial components that play pivotal functional and molecular roles.

**Figure 1.**
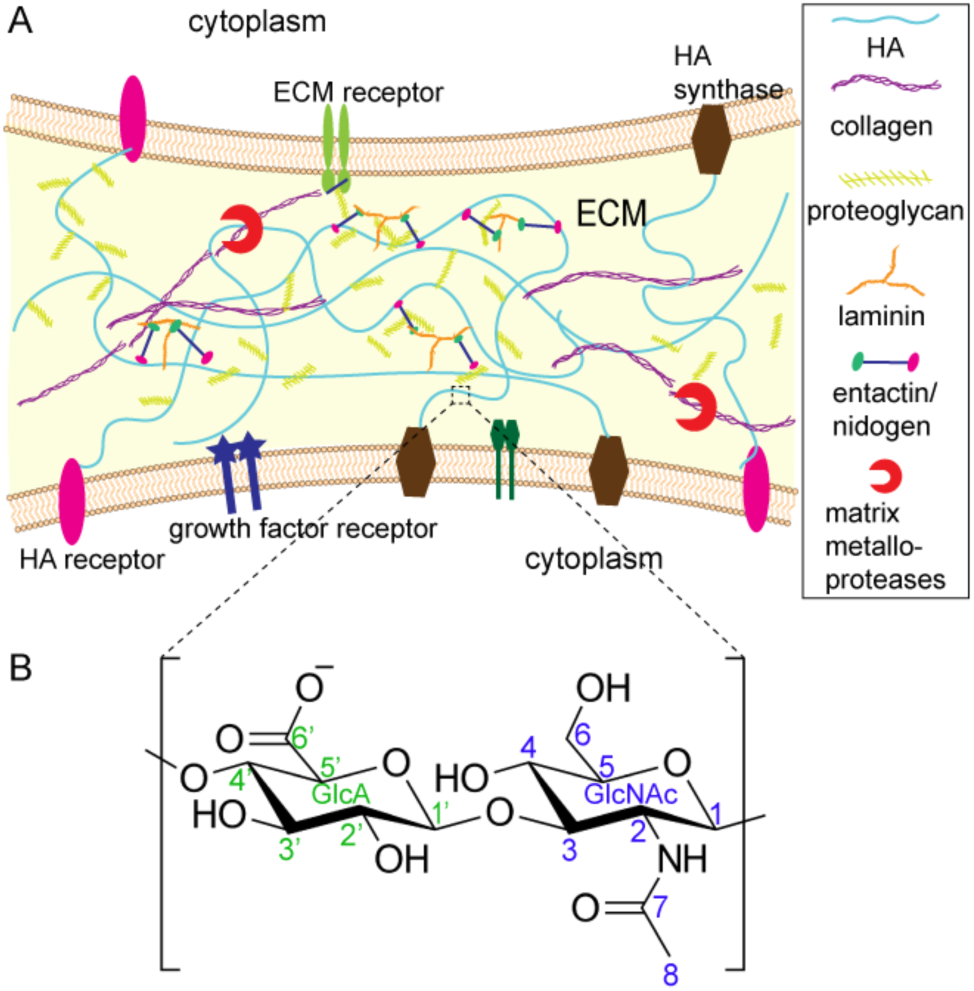
Schematic representation of the ECM and chemical structure of HA. (A) Schematic representation of ECM and its major components. The identity of each component is indicated in the legend. (B) Chemical structure of HA with carbon numbering. Green labels are used for GlcA (glucuronic acid) and blue ones for GlcNAc (*N*-acetyl glucosamine).

HA is a negatively charged, unbranched polysaccharide consisting of glucuronic acid (GlcA) and N-acetylglucosamine (GlcNAc) units (Figure 1B). HA is abundant in connective tissues, such as skin, cartilage, synovial fluid, and brain ECM ^4^. HA serves multiple roles within the ECM. For instance, HA is responsible for the hydration and lubrication of tissues, thanks to its unique structure that enables it to bind and retain large numbers of water molecules^4^. HA can form large, hydrated supramolecular assemblies, which provide cushioning and shock-absorbing properties. This is particularly important in tissues such as brain and cartilage, where HA helps to prevent damage from mechanical stresses ^5^. Additionally, HA offers a framework for interactions with HA-binding proteins and cell surface receptors, influencing many cellular processes. These receptors include CD44, RHAMM, and ICAM-1, with crucial roles in cell signaling, migration, and cell proliferation ^6^.

The molecular understanding of the role of the polysaccharide HA in the ECM lags behind studies of proteinaceous components such as collagen. However, it is clear that especially the size of HA has an intricate effect on its functional properties in physiological processes ^7^. For instance, high molecular weight (HMW) HA exhibits anti-inflammatory properties and it can modulate immune cell behavior, reduce inflammation, and promote tissue regeneration ^8^. In contrast, low molecular weight HA is often associated with pro-inflammatory responses ^9^. Notably, the biological importance of HMW-HA has led to the development of its use in 3D cell culture and biomedical applications. Engineered HMW-HA hydrogels provide a favorable microenvironment for cell growth, differentiation, and tissue formation ^10,11^. It supports cell adhesion, migration, and nutrient exchange, mimicking the physiological conditions *in vivo*^12^. In biomedical applications, HMW-HA is used in tissue engineering, drug delivery, and wound healing therapies due to its biocompatibility, biodegradability, and regenerative properties ^10,13^. Also, for these applications, the MW of the HA is of critical importance.

However, we lack a complete understanding of the molecular underpinnings of how HMW-HA plays these diverse roles, both in context of the multicomponent ECM and in its use in 3D cell culture and other applications. Conformational properties are essential to understand HA function, but the complex contexts and innate structural properties of HA conspire to complicate structural analysis. Structural biology techniques such as X-ray diffraction and liquid-state NMR spectroscopy have been used to investigate the molecular structure and conformations of HA oligosaccharides as well as HA-protein interactions^14–19^. Molecular dynamics simulations have also been performed to probe the structure and dynamics of HA tetrasaccharides ^20–22^. However, these reports mostly cover the structural and dynamic properties of relatively short sections of HA. Our understanding of HMW-HA and HA-based ECM or hydrogels remains limited. This knowledge gap can be attributed to the lack of detailed analysis methods specifically designed for studying these semi-solid and complex samples formed by HMW-HA and its interacting partners. What is needed, is a spectroscopic or structural approach that can analyze hydrated samples of HMW-HA, in presence and absence of a complex mixture of interacting (and non-interacting) components. Here, we deploy advanced solid-state NMR spectroscopy (ssNMR) as a powerful tool to provide this type of information. Traditional solution-state NMR analysis requires rapid tumbling of analyzed molecules, for useful spectra to be obtained. However, this limitation is not applicable when NMR spectroscopy is combined with magic-angle-spinning (MAS), thus enabling the acquisition of high-quality spectra even on solid or semi-solid samples. The use of ssNMR permits the in-depth study of a wide diversity of samples, including non-crystalline substances, human biopsies, hydrogels, tissues, and even whole cells ^23–27^. Modern MAS ssNMR has the potential to provide atomic-resolution insights, even in presence of static or dynamic disorder ^28^, and also information on dynamics of macromolecules through measurements of relaxation and order parameters ^29–31^. Solid-state NMR has proved effective for the study of polysaccharides, in the context of ECM, cell walls, and other contexts ^32–36^. Employing a combination of NMR measurements allows for gaining insights into the structure, dynamics, and functional relationships of polysaccharides. Different NMR methods serve specific purposes, such as probing the structural arrangements of polysaccharide chains, characterizing different components within the polysaccharide matrix, examining molecular connectivities, and determining cell wall structure and molecular interactions ^33,36^. An important feature to note is that ssNMR analysis is not restricted to dry samples, and that it is commonly applied to hydrated (polysaccharide) samples to mimic conditions relevant to biology as well as industrial applications.

One traditional limitation in the application of ssNMR to polysaccharide samples is that the isotopic enrichment common to protein samples is often difficult to achieve. The low abundance of ^13^C nuclei in natural samples limits the utility of more advanced NMR experiments. Here, we produced ^13^C-enriched HMW-HA to perform multidimensional solid-state NMR experiments. Taking advantage of the ^13^C enrichment, we use ssNMR to elucidate the conformational and dynamic properties of the HMW-HA, using a variety of 1D and 2D magic-angle-spinning (MAS). A key asset in these studies is the ability of ssNMR to provide quantitative and site-specific insights into the molecular dynamics of HA, at varying levels of (de)hydration and in complex environments. In a biological (ECM) context the HA is engaged in numerous interactions with other ECM proteins. To examine the impact of a context, we also probe the (hydrated) HA in the context of ECM-mimicking components from the popular Geltrex preparation. Geltrex contains major components of ECM such as laminin, collagen IV, entactin, and heparin sulfate proteoglycans, but is itself devoid of HA. Across this range of conditions, the ssNMR revealed a striking dynamic and structural complexity, affected specific chemical moieties with the HMW HA, seemingly in response to its engagement with the surrounding matrix context.

## 2. Materials and Methods

### 2.1 Production and Purification of ^13^C-labelled HMW-HA

The ^13^C enriched HMW-HA was produced via an optimized protocol, as described in previous work ^37^. Briefly, *Streptococcus equi* subspecies *zooepidemicus* (DSM 20727, obtained from Leibniz Institute DSMZ-German Collection of Microorganisms and Cell Cultures), were sub-cultured and grown by using two percent inoculums containing 3% (w/v) TSB (tryptone soy broth) and 1% glucose. A 500 mL Erlenmeyer flask containing 100 mL of HA production medium, containing (g/100mL): 3 g ^13^C_6_-glucose (Sigma, GmbH); 2 g casein enzyme hydrolysate; 0.3 g yeast extract, 0.2 g NaCl; 0.2 g K_2_HPO_4_; 0.2 g MgSO_4·_7H_2_O, pH 7.0, was used. The inoculated flask was incubated in the shaker incubator at 37 °C and 230 rpm for 24 h. After incubation the broth was diluted with 1 volume of water and clarified by centrifugation at 25000 g for 10 min at 4 °C followed by filtration (0.45 μm syringe filter). The clarified broth containing HA was precipitated with ethanol (1:3 v/v). The precipitated HA was redissolved in MiliQ water and dialyzed against water by using a 3.5 kDa cut off dialysis membrane. Finally, the dialyzed HA was freeze-dried to obtain a powder.

### 2.2 Sample preparation for ssNMR

Dry ^13^C-labeled HMW-HA powder (7 mg) was packed into a thin-wall 3.2-mm zirconia ssNMR rotor (from Bruker Biospin) and an initial series of ssNMR experiments were performed as described below. Next, the sample underwent hydration by the addition of varying amounts of deuterium oxide (D_2_O, Sigma) to achieve specific hydration levels, namely 1:0.5, 1:1, 1:2, 1:5, and 1:8 ratios of HA to D_2_O (w/v). At each hydration step (excluding 1:8 ratio), increasing quantities of D_2_O were introduced to the HMW-HA within the same rotor, and ssNMR measurements were performed. For the 1:8 ratio, the rotor was separately packed with 4 mg of ^13^C-labeled HMW-HA powder and 32 μL of D_2_O (to achieve 1:8 hydration ratio). At each stage, the rotor was kept at room temperature for 24 hours to ensure equilibration before conducting ssNMR measurements. Separately, ^13^C-HMW-HA was mixed with the ECM component mixture Geltrex™ (Thermo, A1413202). Briefly, 4 mg of ^13^C-HMW-HA (dissolved in 400 μl of Milli-Q water) was mixed with 250 μL of Geltrex (concentration ∼15 mg/mL). The Geltrex was used as received. The mixture was incubated at 37 °C for 1 h and subsequently subjected to lyophilization. The entire amount of HMW-HA-ECM lyophilized powder (∼8 mg; of which 4 mg of labeled HA) was then packed into a thin wall 3.2-mm zirconia rotor. Rehydration was achieved by adding D_2_O to the rotor, achieving final HMW-HA-ECM:D_2_O ratios of 1:1.5 (w/v) and 1:4 (w/v), followed by a 24-hour equilibration period at room temperature.

### 2.3 Solid-state NMR spectroscopy

MAS ssNMR experiments were performed using a Bruker AVANCE NEO NMR spectrometer operating at a ^1^H Larmor frequency of 600 MHz (14.1 Tesla) and either a 3.2 mm EFree HCN MAS Probe or a 3.2 mm two-channel HX broadband CPMAS probe from Bruker Biospin. All the spectra were recorded at a temperature setpoint of 277 K and MAS rate of 10 kHz. 1D ^13^C cross polarization (CP), rotor-synchronized refocused insensitive-nuclei enhanced-polarization transfer (INEPT) and direct excitation (DE) experiments were performed to study both the rigidity and mobility of the sample ^30,38^. These 1D experiments used the following parameters: 50 kHz ^13^C nutation frequency (5 μs 90° pulse), 0.01 s acquisition time, 3 s recycle delay and 2 k scans. During acquisition, 83 kHz two-pulse phase modulation (TPPM) ^1^H decoupling was applied ^39^. For 1D ^13^C CP, a 70 %-100 % ramped ^1^H–^13^C CP step was used with contact time set to 500 μs. The 2D ^13^C–^13^C CP-based dipolar-assisted rotational resonance (DARR) ^40^ spectra were recorded using the same CP conditions, with DARR ^13^C-^13^C mixing times of 8 and 80 ms. Here, a 5 μs 90° carbon pulse and 2.5 μs 90° proton pulse were used. TPPM ^1^H decoupling during acquisition was around 83 kHz, recycle delay 3 s and number of scans 16 per datapoint. *J*-coupling-based 2D ^13^C-^13^C spectra, acquired using rotor-synchronized refocused INEPT ^1^H–^13^C transfers and showing ^13^C– ^13^C correlations between highly mobile carbons, were obtained by combining refocused INEPT ^1^H–^13^C combined with 6ms of P9^1^_3_ total through bond correlation spectroscopy (TOBSY) ^13^C-^13^C mixing ^41,42^. Here, 5 μs 90° carbon pulses and 5 μs 90° proton pulses were used. TPPM ^1^H decoupling during acquisition was around 50 kHz, recycle delay set to 3 s and number of scans 128 per datapoint. 2D INEPT-based HETCOR spectra were recorded using ^13^C nutation frequency 50 kHz, ^1^H nutation frequency 83 kHz, 3 s recycle delay, and 256 scans per datapoint, in absence of homonuclear decoupling during the t_1_ evolution. TPPM ^1^H decoupling during acquisition was around 83 kHz. Two-dimensional double-quantum (DQ) correlation spectra were obtained via the INADEQUATE pulse sequence which uses scalar coupling to obtain through-bond information of directly bonded atoms^43^. Carbon 90° and 180° pulse lengths were 5 and 10 μs, with a 2τ spin-echo evolution time for a (π–τ–π/2) spin-echo of 6.24 ms. Around 50 kHz TPPM ^1^H decoupling was applied during both evolution and acquisition times. Number of scans were 64 and 714 complex points were acquired along the indirect dimension.^13^C T_1_ relaxation measurements with direct excitation ^13^C detection were done using the saturation recovery Bruker pulse program (satrect1), using d20 (delay in saturation pulse train) and L20 (number of pulses in saturation pulse train) set to 20 ms and 10, respectively. The recovery delay Tau was varied from 50 μs to 20 s. The ^13^C nutation frequency was 50 kHz, recycle delay 3.5 s, number of scans 1500 and acquisition time 0.01 s. ^13^C T_2_ relaxation measurements with direct excitation ^13^C detection were performed using the Hahn echo pulse program with the following conditions: ^13^C nutation frequency 50 kHz, 3.5 s recycle delay, 2 k scans and 0.01 s acquisition time. The echo time was rotor synchronized and increased from 0 to 6.4 ms. Additionally, ^1^H decoupling was applied 25 kHz during the echo time and 50 kHz during the acquisition. Spectra were acquired with Bruker Topspin, processed with NMRPipe, and analyzed with the CcpNmr Analysis program version 2.4 ^44,45^. The chemical shifts of ^1^H and ^13^C were indirectly referenced to aqueous DSS based on external measurements of the ^13^C signals of adamantane ^46^. Where applicable, during NMR spectra comparison, the vertical scale of the HA-ECM spectra were multiplied by a scaling factor to account for differences in the amount (mass) of HA. MathWorks MatLab and Python were used for plotting graphs and fitting the relaxation data. Peak deconvolution was performed with NMRPipe software using Lorentzian function. The obtained peak areas were then fitted for increasing recovery time or echo time values using a mono-exponential function to extract T_1_ or T_2_ atom specific values. The equations used for T_1_ and T_2_ peak fitting are as follows:

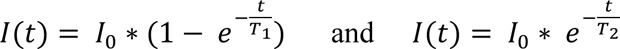

### 2.4 Sample preparation for rheology measurements

Five different samples were prepared for rheology measurements. Among these, three samples had a 1:5 hydration ratio (w/v), and two samples had a 1:10 hydration ratio (w/v) HA:D_2_O. The first sample (1) was composed of HMW-HA mixed with D_2_O at a ratio of 1:5 w/v (20 mg HMW-HA dispersed into 100 μL D_2_O). The second sample (2) consisted of a mixture of 10 mg HMW-HA and 10 mg Geltrex, which is a mix of ECM components derived from murine Engelbreth-Holm-Swarm (EHS) tumors (Thermo Fisher Scientific Inc, Waltham, Massachusetts, U.S.) dispersed into 100 μL D_2_O. The third sample (3) was solely composed of 20 mg Geltrex mixed with 100 μL D_2_O. The fourth sample (4) contained 10 mg HMW-HA mixed with 100 μL D_2_O, while the fifth sample (5) had 10 mg Geltrex mixed with 100 μL D_2_O. All samples were stored at 4 °C for 24 h prior to conducting the rheology measurements. All samples were prepared and measured twice (in duplicate).

### 2.5 Rheology measurements

Dynamic oscillatory shear experiments were carried out with a Physica MCR 300 rheometer (Anton Paar, Germany). The measurements were performed using a 25 mm parallel plate geometry. Samples were prepared and stored at 4 °C for 24 h prior to the measurements. Subsequently, they were placed onto the pre-cooled (at 4 °C) Peltier plate of the rheometer. It was allowed to equilibrate ensuring zero normal stress values prior to the measurement. The linear viscoelastic (LVE) regime was determined using dynamic strain sweeps performed at an angular frequency of 100 rad/s. Consequently, dynamic frequency sweeps were carried out at a chosen strain rate in the LVE regime from 100 rad/s to 0.1 rad/s to delineate the mechanical spectrum of the material.

#### Safety statement

No unexpected or unusually high safety hazards were encountered in this work.

## 3. Results

### 3.1 Atom-specific analysis of hydrated ^13^C-labeled HMW-HA by MAS NMR

To enable a detailed analysis of HMW-HA by ssNMR, we produced ^13^C-labeled HMW-HA using *S. equi* subsp. *zooepidemicus* bacteria^37^. The HA molecules used in this study have a ^13^C enrichment of ∼96 % and an average MW ∼275 kDa^37^. Our initial focus was on elucidating the effect of hydration on the biopolymers, given that the hydration behavior of HA is one of its key functional properties. Thus, we prepared a series of samples with increasing water (D_2_O) content, with a ratio HA:D_2_O ranging from 1:0.5 (w/v) to 1:8 (w/v). The use of D_2_O rather than H_2_O was designed to help suppress the water signal in ^1^H-detected ssNMR. Figure 2 shows 1D and 2D ssNMR spectra for a well-hydrated HMW-HA sample at 1:5 (w/v) ratio. Under this condition, the polymer is highly dynamic, such that it can be detected effectively with the type of scalar-coupling-based NMR techniques common in liquid-state NMR. These INEPT-based techniques work well for flexible molecules but give poor spectra for traditional dry or rigid samples (see below). The ^13^C 1D INEPT spectrum showed good signal to noise ratio (Figure 2A). The resonances were initially assigned based on published solution and ssNMR data ^37,47^, however more peaks were detected compared to the number of HA carbons (Figure 1B). To investigate these unexpected signals in more detail, we performed multidimensional ssNMR experiments. In an INEPT-based 2D ^13^C-^13^C INEPT-TOBSY experiment (Figure 2B) we detected mainly one-bond correlations that allowed complete assignment of all the carbons from both GlcA and GlcNAc moieties of HA. Notably, carbons C1, C2, C4, C5 and C6, belonging to the GlcNAc moiety showed two different forms, which we designate as “a” and “b”. Additional longer-range correlations could be observed, mainly belonging to the GlcNAc moiety (Figure 2B, solid blue lines), including a three-bond correlation involving carbons C6 and C4. We also performed INEPT-based ^1^H-^13^C HETCOR experiments (Figure 2C) to assign the proton resonances. This experiment also revealed multiple conformations for the carbon C6 of GlcNAc moiety. Specifically, four different peaks for this carbon could be detected, indicated as C6a, C6b, C6c and, C6d (chemical shifts in Table S2). Along with C6, also C4 of the GlcNAc moiety showed multiple peaks (three distinct peaks) corresponding to distinct conformations (Figure 2C). The same resonances were not resolved in the ^13^C-^13^C 2D experiment, because of overlap along the carbon dimension (Figure 2B). We complemented the TOBSY data with a ^13^C INADEQUATE (through-bond *J*-mediated) experiment ^43^ that also showed the presence of extra conformations for the carbons C1, C2, C4, C5 and C6 of the GlcNAc moiety (Figure 2D, red solid lines). Thus, we see in this flexible and hydrated HMW-HA a single conformation for the GlcA moieties throughout the polymers, in contrast to a multiplicity of conformations (or dynamics) impacting specific carbons in the GlcNAc.

**Figure 2.**
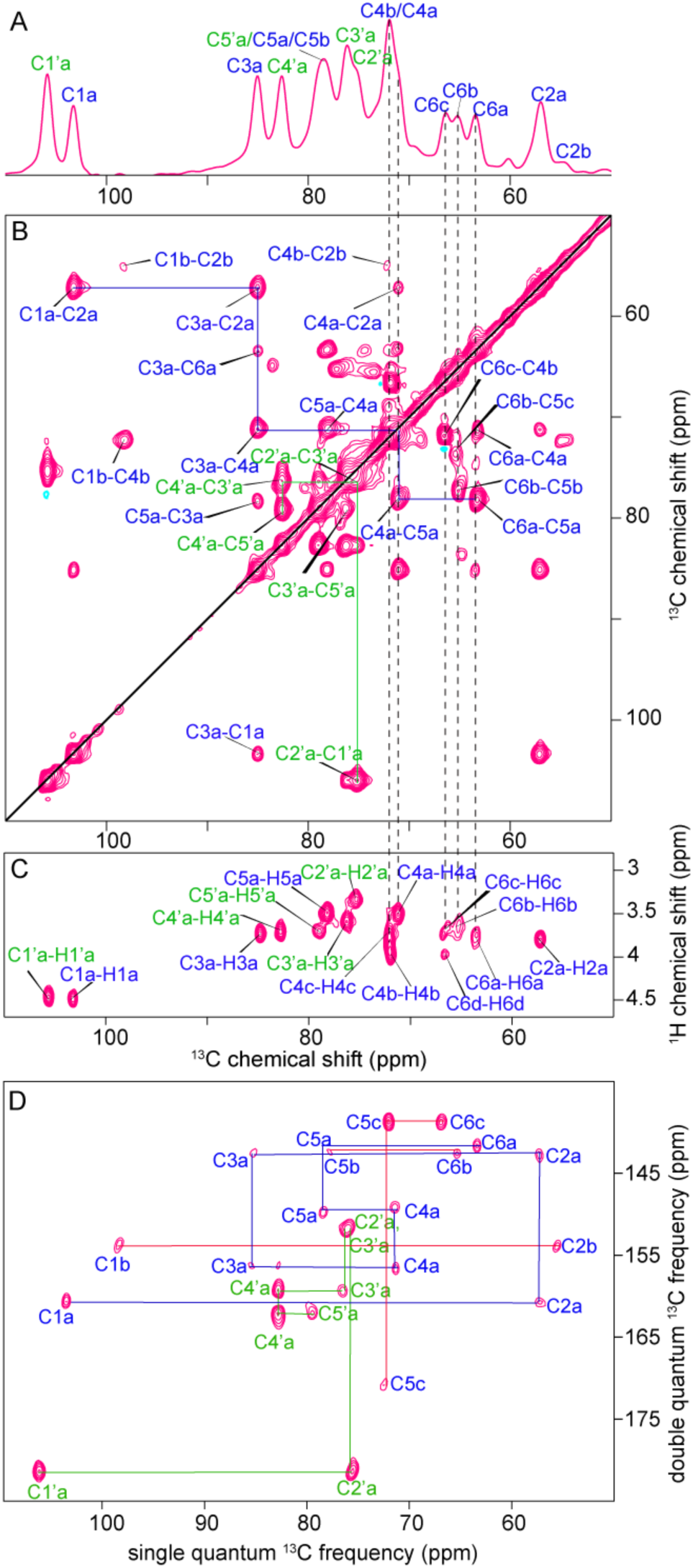
MAS NMR analysis of flexible ^13^C-labeled HMW-HA in the hydrated state. (A) 1D ^13^C INEPT spectrum of HA:D_2_O (1:5 w/v), and (B) 2D ^13^C-^13^C INEPT-TOBSY spectrum on the same sample. The 2D used 6 ms TOBSY mixing time resulting in one-to three-bond correlation peaks. (C) 2D ^1^H-^13^C INEPT-based HETCOR spectrum for HA:D_2_O (1:5 w/v). Vertical dashed lines across the spectra show the presence of multiple signals from the carbons C4 and C6 of GlcNAc indicating the presence of more than one conformation for these two carbons in the hydrated state. (D) 2D ^13^C INADEQUATE spectrum of HA:D_2_O 1:5 (w/v), showing the ring, side chain and anomeric carbons of both moieties. Solid lines show the carbon connectivities for GlcNAc (blue) and GlcA (green). Red lines indicate extra conformations of GlcNAc moiety. The carbonyl and methyl peaks are visible in the full range 2D ^13^C INADEQUATE spectrum, which is reported in Figure S1.

### 3.2 Global and local structural changes as a function of hydration

To gain more detailed insights into the conformational features of HMW-HA and the impact of HA hydration on the conformation, we applied MAS ssNMR spectroscopy to ^13^C-labelled HMW-HA at a range of hydration levels (from 1:0.5 to 1:8 hydration ratio, w/v). ^13^C spectra acquired with the INEPT technique (Figure 3A and Figure S2), gave a good signal intensity and spectral resolution for the most hydrated states (1:5 and 1:8 hydration ratio). Notably, upon further increasing hydration from 1:5 to 1:8, no significant differences were observed in terms of structural conformations (Figure 3A and Figure S2). However, as we reduced the hydration level, the observed signal intensities dropped dramatically. A notable feature in this behavior is that different peaks responded differently to the reduced hydration level, revealing an apparent preferential hydration of specific parts of the HA polymer. In the black spectrum in Figure 3A, showing a 1:1 hydration level (by weight), it is striking that the main HA peaks were largely lost, but signals from the ‘minor’ sub-states remained (extra peaks corresponding to secondary forms of C2, C4 and C6). As before, these peaks belong to the GlcNAc moiety, suggesting that it is preferentially engaged with fluid water in these semi-dry sample conditions, potentially forming local pockets of increased hydration. Yet, in these medium- to low-hydration conditions (1:1 ratio), the characteristic flexibility of most of the HA polymer chain is clearly lost.

**Figure 3.**
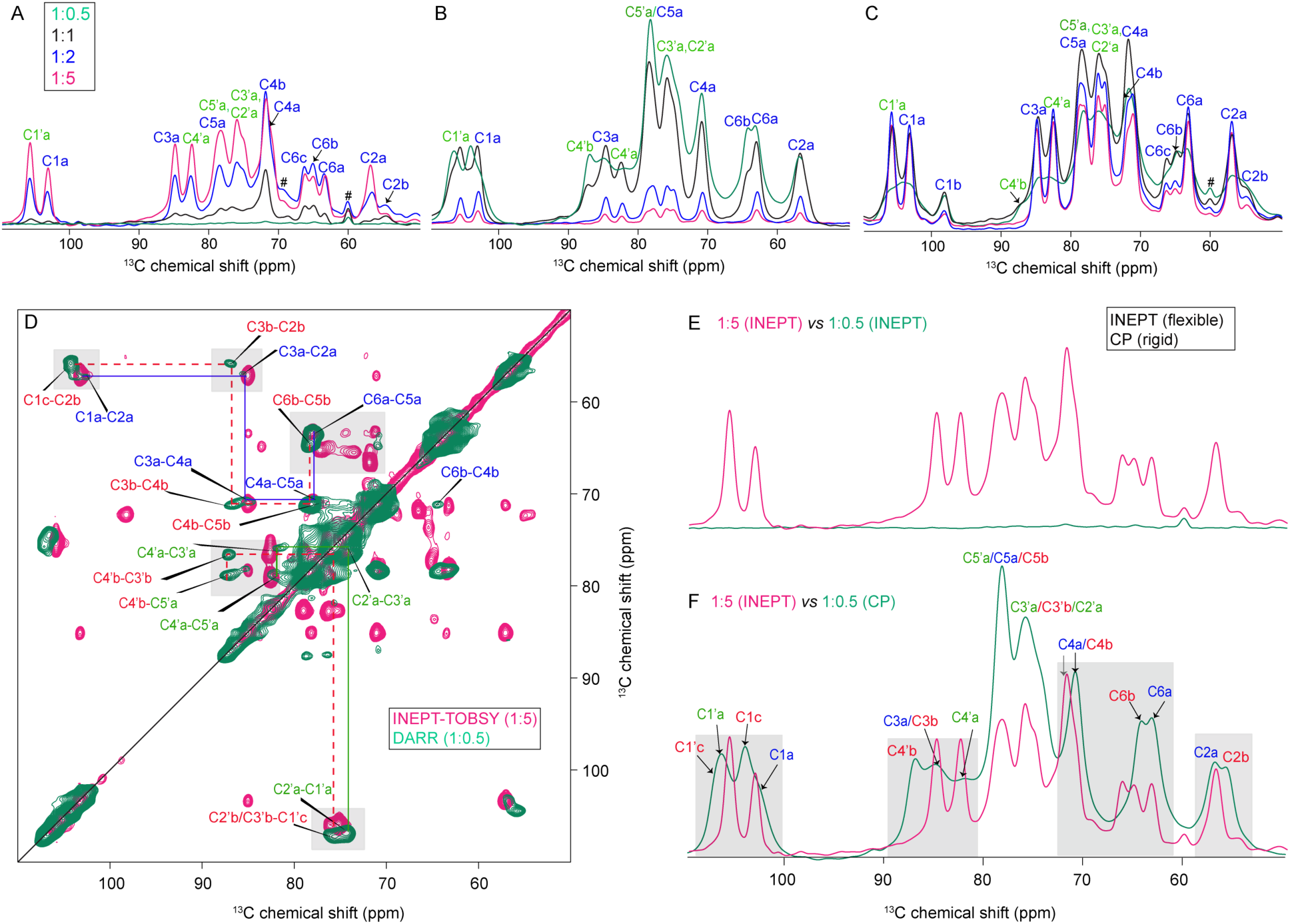
Hydration-dependent site-specific changes in HMW-HA. (A) INEPT, (B) CP, and (C) DE ^13^C 1D ssNMR spectra of HMW-HA at different hydration levels (green 1:0.5, black 1:1, blue 1:2, pink 1:5 hydration ratio). Hash signs (#) indicate signals that derive from an impurity in the sample. The full range spectra are reported in Figure S5. (D) Overlay of 2D DARR (low hydration, green) and INEPT-TOBSY (high hydration, pink) spectra. (E) Overlaid 1D ^13^C CP and INEPT spectra of HA (1:5 hydration ratio, w/v). Low CP signal and high INEPT signal indicated high flexibility of HA at this hydration level. (F) Spectral comparison of low (1:0.5, green) and high hydration (1:5, pink) of two different hydration states of the same sample as in panel D: flexible signals from the high hydration (INEPT) were compared to rigid signals at low hydration (CP), showing similarities and differences. Gray boxes in D and F mark glycosidic carbons and side chain carbons which experienced the most prominent hydration effect. Red dashed lines in D show the carbon-carbon connectivity map for extra forms in both moieties (prime sign for GlcA and without prime is GlcNAc). Signals from GlcA carbons are indicated with green labels and a prime sign, and signals from GlcNAc carbons are indicated with blue labels.

Under such conditions, 1D ^13^C cross-polarization (CP) ssNMR is a classic approach for studying immobilized (semi-)solids, as this ssNMR technique requires a lack of fast isotropic motion (Figure 3B). To interpret these CP spectra, we performed both 1D and 2D measurements (see below). At high hydration levels, the HA is sufficiently flexible that it cannot be detected in the CP based experiments. When the hydration level was decreased, the overall intensity and quality of the CP spectra improved, complementing the INEPT analyses above and reflecting the hydration-associated changes in HA flexibility. Several interesting changes were observed. For instance, at the lowest hydration level specific carbons from the GlcA moiety showed extra forms, i.e. carbon C4’ appeared at two distinct positions (labeled as ‘a’ and ‘b’, at 81.9 ppm and at around 87.7 ppm respectively, Figure 3B). Similarly, the anomeric carbon C1’ also showed two different forms, at 106.8 ppm and 107.2 ppm (Table S2 and Figure 3B, green and black spectra). The anomeric carbon C1 and ring carbons C2 and C6 from the GlcNAc moiety also showed extra forms, assigned as form ‘a’ and ‘b’. Though these forms are not always clearly distinguishable in the 1D spectra, they became clearly visible in 2D ^13^C-^13^C CP-DARR spectra (Figure 3D and S3, S4). Interestingly, at increased hydration levels, these ‘extra’ signals were less prominent or not detectable, indicating that hydration renders the HA’s behavior (and conformation) more uniform. As discussed above, the combination of INEPT- and CP-based spectra permits a qualitative analysis of changes in flexibility/rigidity. This approach is sometimes discussed as ‘polarization transfer’ (PT) or ‘dynamics-based spectral editing’ (DYSE) analysis ^30,48^. However, these techniques do not permit a (semi)quantitative analysis of the population sizes of molecular or conformational states, since the peak intensities depend on both the number of detected molecules and their motion. For a more quantitative analysis we turned to single pulse excitation (SPE) or direct excitation (DE) experiments, where a 90° pulse is applied on the nucleus of interest (e.g. ^13^C)^30^. In this type of experiment, the integrated signal intensity is considered a more reliable measure of occupancy or population size. Indeed, we observed that the *overall* spectral signal intensity was largely independent of the hydration level, as it reflects the overall amount of HA, which did not change (Figure 3C). However, interesting hydration-dependent changes were observed *within* the spectra, illuminating hydration-dependent changes in structure. At the least hydrated state (1:0.5 HA: D_2_O), the peaks became very broad, due to a static heterogeneity absent at the more hydrated states. It is also notable that peaks reflecting ‘minor’ conformations varied in intensity as a function of hydration. For instance, the peaks marked as C1b, C6b and C6c were present at high hydration levels, but increased in relative intensity as the hydration level was reduced. Notably, some peaks were seen in these spectra that were absent in the CP spectra (C6c and C4b), indicating that they must reflect dynamic conformers of HA.

### 3.3 Hydration-dependent site-specific changes observed in two different hydration states

In Figure 3D-F we compare the ssNMR spectra for a low- and high-hydration HA state, to illustrate the specific hydration-related changes. The former (low hydrated state) is shown via its CP-based 1D and 2D spectra, and the latter from INEPT-based ssNMR data. Strikingly, the glycosidic linkages showed prominent changes, with a downfield shift of about 1 ppm from the highest to the lowest hydration level (C1’a-C3a and C1a-C4’a, solid boxes in Figure 3D and F, and Figure 4A and 4B). As seen in the 1D spectra, the lowest hydration state exhibited multiple conformations (Figure 3D, green spectrum). Most of the carbons showed more than one peak, indicating multiple coexisting conformations in both the moieties (red dashed lines indicating connectivities for the extra conformations are shown in Figure 3D, green spectrum). However, for the highest hydration level multiple conformations were only observed in carbon C2, C4 and C6 of the GlcNAc. These multiple conformations are labeled as C2a, C2b; C4a, C4b, C4c; and C6a, C6b, C6c, C6d (Figure 3F, pink spectrum, and Figure 4D and 4E).

**Figure 4.**
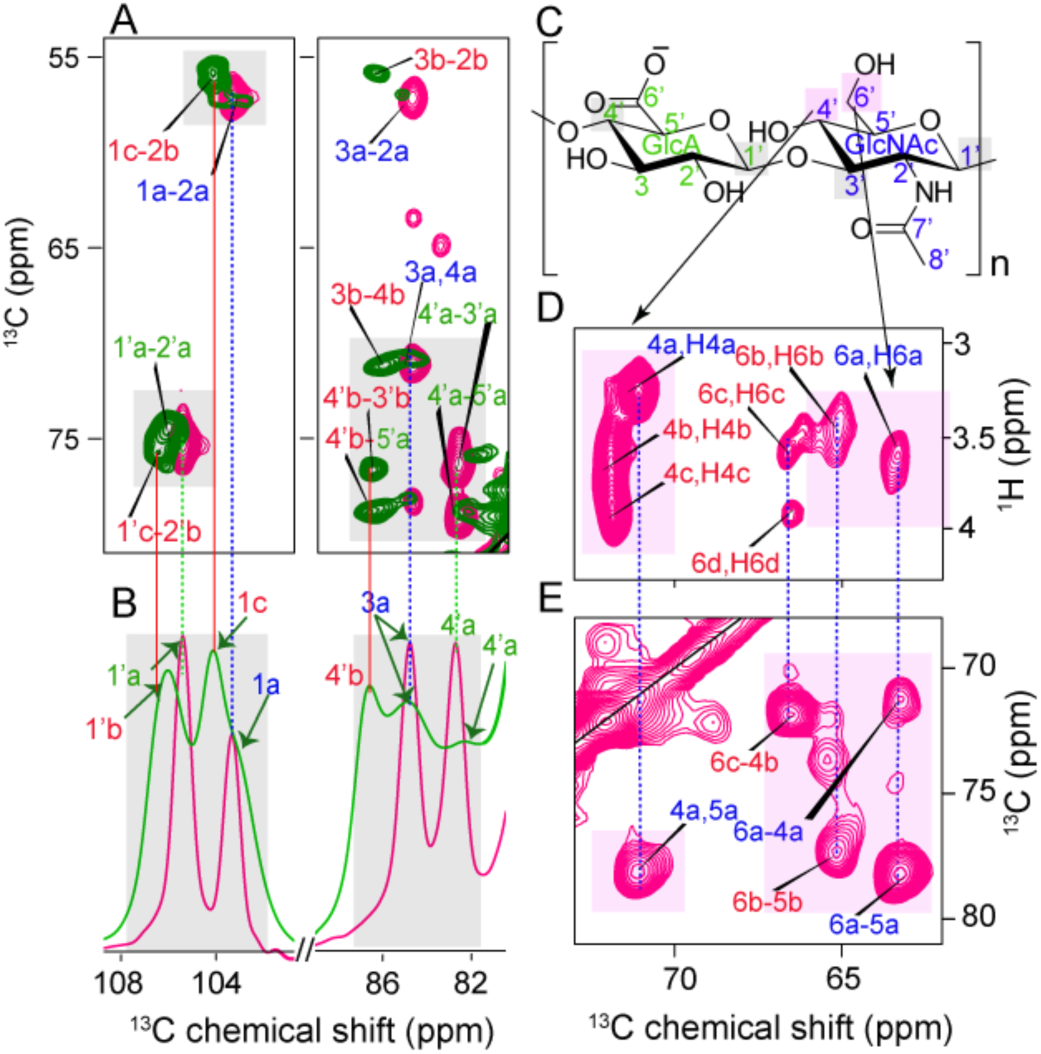
Hydration-induced multiple structural conformations within the GlcNAc moiety. (A) Overlay of 2D DARR (green, lowest hydration state) and 2D INEPT-TOBSY (pink, highest hydration state). (B) 1D CP (green, lowest hydration state) and 1D INEPT (pink, highest hydration state). (C-E) For the highest hydration state, the most affected peaks can be seen in the 2D ^1^H-^13^C INEPT-HETCOR and INEPT-TOBSY spectra.

Thus, we observed by ssNMR that dehydrated HA was conformationally heterogeneous and clearly distinct from the highly flexible, but also homogeneous, native-like hydrated HA polymers. In the latter, structural heterogeneity was limited to specific parts of the GlcNAc moiety, which notably also seemed involved in preferential hydration at intermediate hydration levels.

### 3.4 Structure and dynamics of HMW-HA in an ECM-like environment

The above illustrates the potential for combining 1D and 2D MAS NMR with ^13^C-enrichment to study in detail the behavior of (hydrated) HMW-HA polymers. Next, we set out to deploy these methods to study HMW-HA in a more complex and biologically relevant system, designed to replicate features of native ECM conditions. We used a biomimicking ECM mixture called Geltrex^TM^ matrix which has the same major components (laminin, entactin, collagen type IV, and heparin sulphate proteoglycans) as the biological ECM, except for the crucial component HA. This type of mixture is commonly used to prepare ECM-like hydrogels for cell culture and other applications ^49,50^. The ^13^C-labeled HMW-HA was mixed with ECM and a HA-ECM hydrogel was prepared. To understand the mechanical properties of this hydrogel, we performed rheological studies on HMW-HA and the HMW-HA-ECM-like complex. As shown in Figure 5A, the pure HMW-HA samples exhibited liquid behavior. A crossover of the loss modulus (G”) over the storage modulus (G’) occurred at a higher frequency in the more hydrated conditions. Naturally, this is expected at lower polymer concentrations because the chains can relax much faster. In contrast, the HMW-HA-free ECM (Geltrex) displayed G’ values higher than G” over the entire frequency range, which is indicative of solid-like behavior. Looking at the reconstituted HMW-HA-ECM complex, we observed a behavior that is very distinct from pure HMW-HA, representing a solid response more like the non-HA ECM components in Geltrex. To compare the samples, the tan δ values were plotted as a function of the angular frequency (Figure 5B). The dissipation factor or loss angle, tan δ, is the ratio of G” to G’ indicative of the physical nature of a material. A tan δ value greater than 1 implies a liquid-like material, whereas less than 1 indicates a solid-like material. We can observe that the tan δ values clearly demonstrate a distinction between the HA-free GelTrex mixture and HMW-HA-ECM complex hydrogels. The latter exhibited higher tan δ values, indicative of more liquid-like behavior. This can be attributed to the lower rigidity of HA as compared to the other components of the ECM. Therefore, incorporating HA leads to hydrogels whose stiffness is largely dictated by the non-HA components, although the overall flexibility of the obtained ECM hydrogel is increased by the presence of the HA.

**Figure 5.**
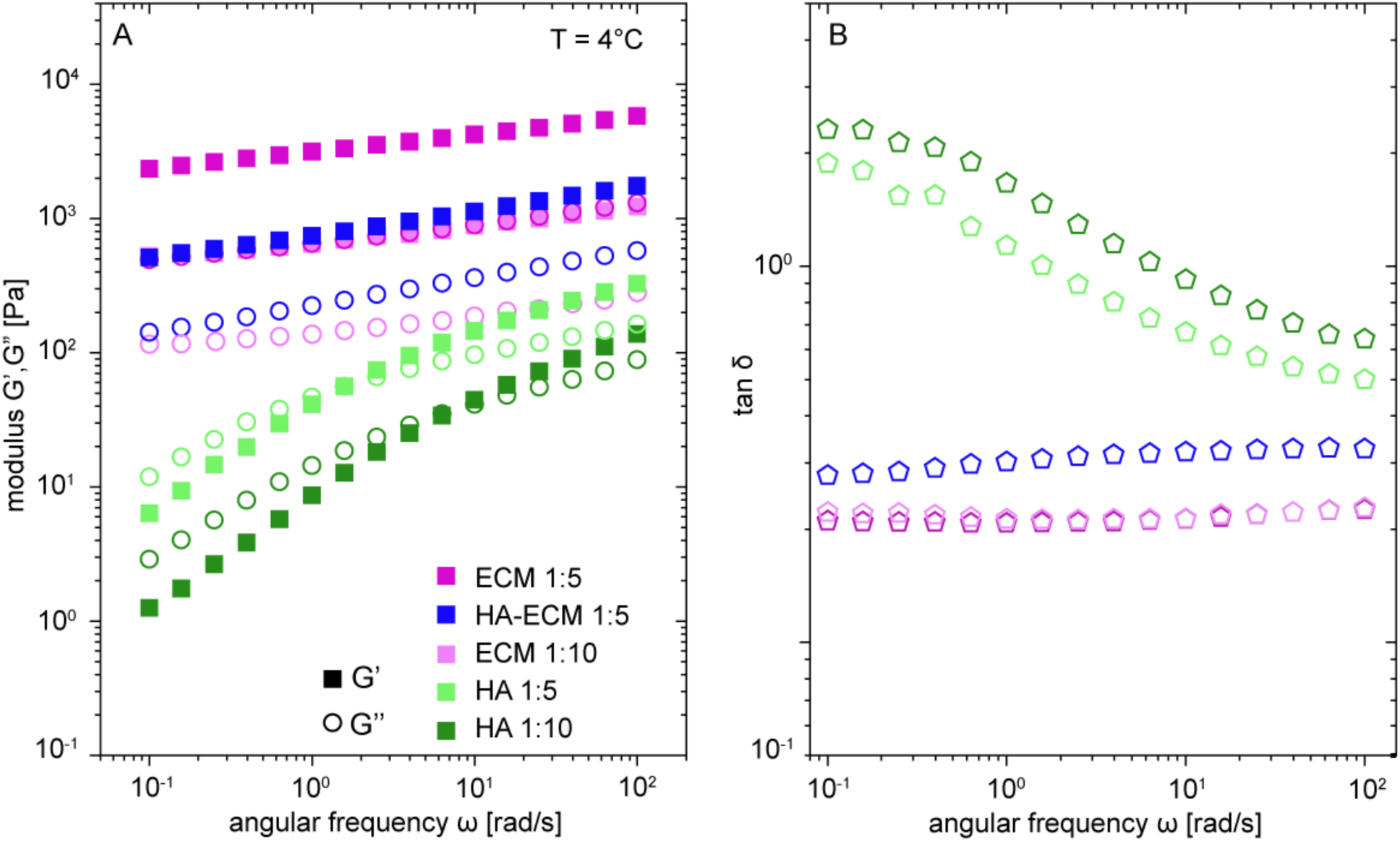
Rheological analysis of hydrogels formed by HMW-HA and ECM with different hydration levels. (A, B) Dynamic frequency sweep data for HMW-HA alone and HMW-HA-ECM complex hydrogel compared with their individual components (see legend) plotted as (A) elastic modulus G′ and loss modulus G″ vs. angular frequency ω. and (B) tan δ vs. angular frequency ω.

Thanks to the selective ^13^C enrichment of the HA, we could now use ^13^C MAS NMR to study the behavior of the HMW-HA polymer in this complex ECM-like context, with its distinct mechanical properties. First, we studied the HA-ECM mixture in the presence of a four-fold excess of D_2_O (1:4 hydration ratio, w/v) using the same NMR experiments as for HMW-HA alone. In the INEPT experiment, we observed that HA in this ECM-like environment is highly flexible (Figure 6C and Figure S6A). Interestingly, even though the hydration level here was slightly lower than in the 1:5 hydration level used for pure HA (due to experimental limitations), we observed a higher INEPT signal for HA in this ECM context. Especially carbons from the GlcNAc moiety showed increased INEPT intensity, whereas the DE experiments showed no noticeable differences between the samples (Figure S6B). Thus, HA in an ECM-like environment, specific carbons from the GlcNAc become more flexible in comparison to HMW-HA alone.

**Figure 6.**
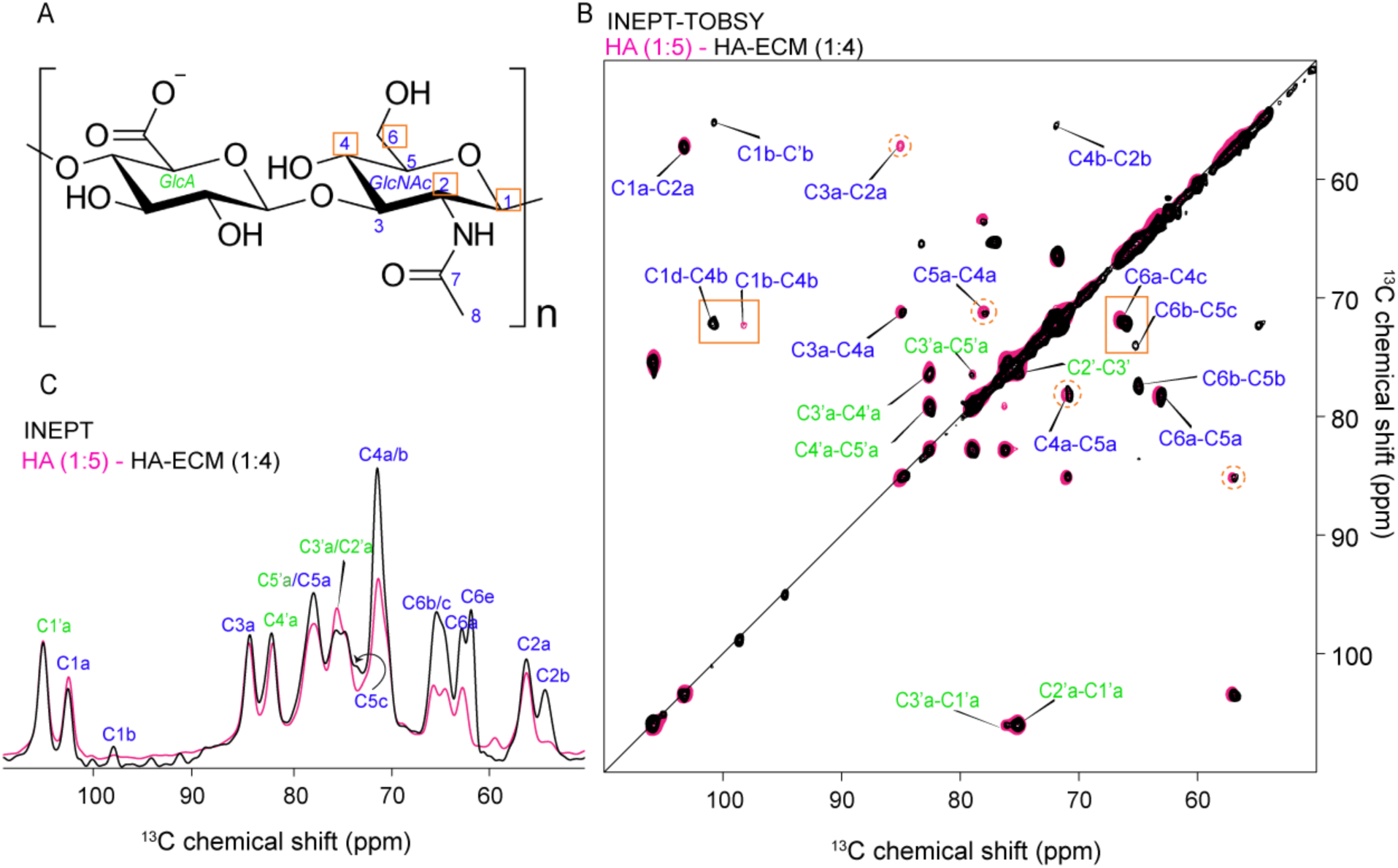
Effects of a hydrated ECM-mimicking context on HA. (A) Chemical structure of HA highlighting the most affected carbons due to the ECM mimicking context (orange boxes). (B) Overlayed 2D ^13^C-^13^C INEPT-TOBSY spectra of HA (pink) and HA-ECM (black). (C) 1D ^13^C INEPT spectral region of HA (pink) with HA-ECM (black) (full spectra in Figure S6). Orange boxes highlights carbons which show difference in the peak intensity, chemical shift, and peak multiplicity in the HA-ECM sample. The dashed orange circles are indicating carbons which show peak asymmetry.

2D ^13^C-^13^C INEPT-TOBSY and ^1^H-^13^C INEPT HETCOR experiments (Figure 6B and Figure S7) revealed more detailed changes. Some of the GlcNAc carbons, such as C1 and C6, showed chemical shift perturbations compared to HA (without ECM), with the former showing the largest chemical shift difference (∼2.5 ppm; marked C1d and Table S3). In contrast, peaks from GlcA remained unchanged in the HA-ECM sample. Thus, the ssNMR analysis revealed that specifically the GlcNAc moiety was showing significant conformational changes with and without ECM components. Especially, carbon C6 and C4 showed the most prominent changes as a function of hydration and in interaction with the ECM environment. Notably, it has been reported in various studies that these sites may be crucial in stabilizing the interactions with HA binding receptors (see _below) 16,17,51._

### 3.5 Dynamical changes in HA as a function of context

The comparison of NMR spectra measured with different polarization transfer techniques pointed to significant differences in dynamics in HA as a function of hydration and ECM-like context. To probe changes in HA dynamics in more detail, we also measured the atom-specific NMR relaxation properties of the ^13^C-labeled HA. For HA and HA-ECM hydrogels with different hydration levels we performed ^13^C T_2_ (spin-spin) and T_1_ (spin-lattice) relaxation measurements, which can reveal different modes of molecular motions on the ps-ns or ns-μs timescale, respectively (Figure S8, Figure S9 and Figure S10). Consistent with the INEPT intensity changes, increasing the hydration led to increases in T_2_ values from less than 2ms to 12ms. All carbon atoms experienced this increase, indicating an overall increase in polymer chain flexibility and reduced (faster) correlation time of molecular motion, with the chain undergoing dynamics on the ns-μs timescale. However, larger effects were observed for the GlcNAc moiety, especially for the carbons C1b, C2b, C6a and C6b, suggesting increased local motion in that time regime. The T_1_ relaxation time was observed to shorten with increasing hydration, which would be expected for a polymer with increasing dynamics at, or approaching, the ns timescale. We interpret these dynamics results to indicate a HA chain that is flexible, meaning that individual atoms (or chemical groups) in the chain are able to access such (ns timescale) motions. This contrasts with the behavior of (folded) proteins but also polysaccharides in higher-order structures (e.g. cellulose), where such chain flexibility is suppressed or absent. The lowest T_1_ values (around 0.1 s) were observed for the carbons C1b, C2b, C6a, C6b and C6c, again specific to GlcNAc, suggesting increased dynamics in GlcNAc relative to the overall HA chain. Furthermore, the comparison of T_2_ and T_1_ values between HA alone (1:5 hydration) and HA-ECM (1:4) showed mostly similar results. This is somewhat surprising given the latter samples’ slightly lower hydration state which would be expected to reduce HA motion, based on the hydration studies above. Therefore, the ECM components do not significantly reduce the general (backbone) dynamics of the HA chains, despite their presence within a more rigid matrix as seen by rheology. Nonetheless, modest relaxation differences were observed to affect the GlcNAc carbons C1, C2 and C6. These showed slightly lower T_2_ values in the presence of the GelTrex components, which presumably reflect interactions of those carbon atoms with the surrounding ECM components.

## 4 Discussion

To gain a comprehensive understanding of the structure and conformation of HMW-HA without and within an ECM-like environment, we combined ^13^C enrichment with multidimensional solid-state NMR experiments. Our findings revealed that the hydration level and surrounding components, such as the ECM environment, significantly impact the overall motion of the HMW-HA chain. These differences highlight the importance of studying biomolecular polymers like HA in a hydrated form and of high molecular weight, and preferably native-like condition is crucial, given that dry (or e.g. crystalline) conditions and short oligomers may not recapitulate the conformational features relevant to their biological function ^14,20,22^. Our studies indicated that especially the GlcNAc moiety was highly responsive to both ECM interactions and hydration-related effects. Surprisingly, no detectable changes were seen for the GlcA moiety, at least under high-hydration conditions. By utilizing multidimensional NMR, we identified multiple conformations of HA that are occupied in the hydrated state. Through 2D experiments, we assigned the carbons from both the GlcA and GlcNAc moieties of HA^37,47^. Furthermore, the use of INEPT-based ^1^H-^13^C HETCOR experiments facilitated the assignment of proton resonances and unveiled multiple conformations for the carbon atoms C6 and C4 of the GlcNAc moiety. These discoveries emphasize the flexibility and existence of various conformations of HA under different hydration conditions, revealing that this variability in structure and dynamics is particular to the GlcNAc of HA.

HA is well-known for its good hydration capacity, enabling very high amounts of water to be engaged by the HA polymers^4,10^. We explored the effects of hydration on the structure and dynamics of HA. Comparisons between HA alone and HA within the ECM hydrogel showcased intriguing spectral changes at a specific hydration level, suggesting an influence of the ECM on HA’s motion and ability to engage with the surrounding water molecules. HA within the ECM-like environment exhibited greater flexibility compared to HA without ECM, considering comparable hydration levels. These dynamics were detected in an atom-specific manner through analysis of ssNMR spectra based on polarization transfer techniques sensitive to motion, as well as explicit NMR relaxation measurements. T_1_ and T_2_ relaxation experiments demonstrated that higher hydration levels increased the overall dynamics (i.e., chain fluctuations or motions on the ns timescale) of the HA chain, while atomic specific effects were observed that mostly affected the GlcNAc moiety. In NMR studies of (small) globular molecules one commonly considers tumbling in solution as a dominant mode of motion in analyzing solution-state NMR relaxation data. Here, these large macromolecules, entangled in densely packed conditions, are unable to undergo overall tumbling, but we rather attribute these motions to chain flexibility, described by the correlation time of relevant dynamics. The relatively high dynamics of polysaccharides in the ECM, compared to e.g. collagen and other protein components, have been noted in previous (NMR) studies^34,35,52^. Here, the use of ^13^C-labeling enabled the detection of interesting variations in local motion within HA. Lower T_2_ values were observed for C1, C2 and C6 carbons, indicating local *decreases* in dynamics on the ns-μs timescale relative to HA alone, which we attribute to the effects of interactions with the ECM components (from GelTrex). These findings appeared to diverge from the results of the INEPT data, which revealed increased intensity (normalized for the mass of HA present in the samples) for the same carbon atoms (C2, C4, and C6). One challenge is that both types of experiments (DE and INEPT) represent an averaged representation of the sample as a whole, which is expected to contain molecules or molecular segments with varying mobility. This is especially true in ECM or hydrogel contexts, where cross-links (whether physical, chemical or otherwise) would be expected to locally reduce chain- or molecular motion. In this context, it is essential to recognize that the INEPT experiment is a non-quantitative experiment that exclusively reflects the most flexible segments or molecules within the sample, with slow-moving or rigid groups even being undetected by INEPT measurements^30^. Thus, the peak intensities in the INEPT spectra depend on both the number of detected molecules (with sufficiently long T_2_ values) *and* on the value of the T_2_ relaxation for the molecules (longer T_2_ gives more intense peaks). In contrast, the T_2_ data we analyzed stemmed from direct excitation (DE) experiments, which should reflect the whole sample. Indeed, the DE-detected spectra are often considered (semi-)quantitative, as they are less sensitive to local dynamics. An important caveat is that the quantitative interpretation can be imperfect even for DE ssNMR measurements, given the need to account for components with long T_1_ values, which are underrepresented if the recycle delay is insufficiently long, possible difficulties in fitting (broad) signals in presence of baseline distortions, and the effect of receiver dead time, resulting in under-estimating signals with fast T_2_ relaxation. With all these considerations in mind, we rationalize the seemingly contradicting results from the INEPT and DE data as follows. The latter reveals an overall decrease in HA dynamics in the sample, which we attribute to the effect of local reductions in dynamics under ECM-mimicking conditions. This is consistent with the effect of cross-links between the HA and the ECM, which would (locally) impede the natural motion of the HA chains. The increased INEPT signals can be explained by sub-population of HA segments distinct from these crosslinks experiencing faster dynamics, leading to longer T_2_, resulting in stronger INEPT peaks, which we will discuss more below. Thus, we interpret these combined data to indicate localized reductions in HA dynamics (on the relevant timescales), where interactions with ECM components arise, whilst other HA parts are not engaged in such interactions and able to stay flexible.

Indeed, we noted a surprisingly *enhanced* dynamics of HA when mixed with ECM components (Figure 6). Given the noted connection between hydration and HA flexibility, we attribute this effect to the ability of HA (related to other ECM components) to undergo preferential hydration. Thus, when a mixture of HA and other ECM components (e.g., collagen) are exposed to a given amount of water, more of this water becomes engaged with the HA than with other ECM components^52,53^. While this facilitates the enhanced mobility of the HA seen by NMR, it also offers the intriguing implication that HA addition effectively decreases the apparent hydration of other ECM components. Further (ssNMR) studies may be warranted to investigate this potent crowding effect, both in its relevance to only biological ECMs but also in HA-based hydrogels for biomedical applications^54^.

Taken together our ssNMR spectra clearly showed the importance of hydration for the HA polymers to explore their dynamic and flexible behavior. A striking finding across all our NMR analyses is the observation that specific carbons in GlcNAc were most sensitive to changes in the environment and interactions. Spectral multiplicity selectively affected C4 and C6 of the GlcNAc moiety, seen both as a function of hydration and upon ECM interactions. The GlcNAc moiety, specifically carbons C1, C2, and C6, also displayed higher T_2_ values and lower T_1_ values compared to other HA parts, indicating increased dynamics. A particular involvement of GlcNAc may at first glance be anticipated. However, one might have expected a special role for the N-acetyl group of GlcNAc moiety, which is not what we observed. The ^13^C and ^1^H resonances of that group proved notably less sensitive to environmental changes than the C4 and C6 hydroxyl groups (and nearby carbons) from the same moiety. This led us to further examine previous reports on the roles of these carbons in dictating the structure and interactions of HA. The formation of an intramolecular hydrogen bond between the hydrogen of the hydroxyl group at C4 of GlcNAc and the ring oxygen (O5) of GlcA is thought to play a crucial role in stabilizing the structure of HA. Figure S11A shows the published structure of the HA-binding domain (HABD) of CD44 bound to a HA oligosaccharide ^16,17^. The inset shows the conformation of the HA, and the presence of a C4-OH hydrogen bond between GlcNAc and GlcA in the bound carbohydrate. Similarly, this hydrogen bond is present in the integrative HA model shown in Figure S11B, based on MD and solution NMR studies ^55^. This hydrogen-bonding interaction is thought to maintain HA’s overall architecture, and it is retained in the bound and unbound HA. The variability of the C4 signals in our NMR studies suggest a certain degree of flexibility or heterogeneity in this stabilizing interaction, at least in our HMW-HA polymers. The observed heterogeneity in the C6 site is also intriguing. Figure S11A shows that this hydroxyl group is implicated in HA recognition by the CD44 HABD. The oxygen attached to C6 is forming a hydrogen bond with the hydrogen of the hydroxyl group of the Tyr 109 (Figure S11A inset and Figure S12) ^16,17^. This part of HA has also been implicated in HA interactions by serum-derived HA associated proteins (SHAPs). A C-terminal Asp of SHAPs binds covalently with the C6-hydroxyl group of GlcNAc^51^. The importance of this interaction was illustrated by the fact that elimination of the SHAP-HA binding complex led to the severe female mice infertility ^56^. An intriguing interpretation of the observed multiplicity of the GlcNAc C4 and C6 signals is that this HA moiety is able to occupy several different conformations, depending on intra- and inter-molecular interactions as well as solvent effects. HA-binding proteins may then recognize this flexible HA region, reflecting a type of conformational selection in the recognition of (HA) substrate and HA-binding proteins^57^.

## 5 Conclusion

Overall, our study provides insights into the conformational non-HAns of HA under varying hydration conditions and within an ECM-like environment. The results obtained through MAS NMR, supported by rheological measurements, provide atom-resolution insights into the flexibility of HA and its capacity to undergo structural changes upon hydration and supramolecular interactions, depending on the environment. Multiple conformations in highly localized and unexpected parts of the HA polymer responded to hydration and ECM interactions, potentially reflecting hinge-type regions in the polymer that are important to its mechanical properties and recognition by HA binding proteins. The combination of ^13^C-enrichment of HMW-HA and multidimensional MAS NMR is a powerful approach to understand HA’s behavior and interactions. We foresee this approach to be instrumental for a better understanding the diverse biological functions of HA, but also for molecular studies of HA-based nanoparticles and hydrogels with important biomedical and pharmaceutical applications in various industries.

## Supporting Information

Tables with detailed solid-state NMR experimental parameters and ^1^H and ^13^C chemical shift assignments; additional 2D and 1D solid-state NMR spectra; T_1_ and T_2_ relaxation plots; illustrations of molecular structural models of HA in its bound and unbound states.

## Acknowledgment

This work was supported by financial support from the Zernike Institute for Advanced Materials, University of Groningen, including funding from the Bonus Incentive Scheme of the Dutch Ministry for Education, Culture and Science (OCW).

