## Supporting information for "Resolving atomic-level dynamics and interactions of high molecular weight hyaluronic acid by multidimensional solid-state NMR"

**Table S1: Experimental conditions of ssNMR experiments:** Abbreviations: NS, number of scans per $t_1$ point; RD, recycle delay; TPPM, $^1$H decoupling power during evolution and acquisition using the two-pulse phase modulation scheme; Mixing, $^{13}$C-$^{13}$C or $^1$H-$^1$H mixing time (ms); $t_1$ evol., maximum $t_1$ evolution time expressed in number of $t_1$ points (real+imaginary) x $t_1$ increment time. The experiments were conducted at a temperature of 277K using a magic angle spinning (MAS) of 10kHz.

| Figure | Sample | Experiment | NS | RD (s) | TPPM (kHz) | $t_1$ evol. (□s) | Mixing (ms) |
|---|---|---|---|---|---|---|---|
| 2A, 3A, 3E (pink), 3F (pink), 4B (pink), 6C (pink), S2A, S5C, S6A (pink) | HMW-HA (1:0.5, 1:1, 1;2, 1:5 hydration state) | $^1$H-$^{13}$C INEPT | 2048 | 3 | 83 | N/A | N/A |
| 3B, 3E (green), 3F (green), 4B (green), S5A | HMW-HA (1:0.5, 1:1, 1;2, 1:5 hydration state) | $^1$H-$^{13}$C CP | 2048 | 3 | 83 | N/A | N/A |
| 3C, S5B, S6B (pink) | HMW-HA (1:0.5, 1:1, 1;2, 1:5 hydration state) | $^{13}$C DE | 2048 | 3 | 83 | N/A | N/A |
| 6C (black), S6A (black) | HMW-HA-ECM (1:4 hydration state) | $^1$H-$^{13}$C INEPT | 2048 | 3 | 83 | N/A | N/A |
| S6B (black) | HMW-HA-ECM (1:4 hydration state) | $^{13}$C DE | 2024 | 3 | 83 | N/A | N/A |
| 2C, 4D, S7 (teal) | HMW-HA (1:3, 1:5 hydration state) | 2D $^1$H-$^{13}$C INEPT-HETCOR | 256 | 3 | 83 | 108x185.15 | N/A |
| S7 (mauve) | HMW-HA-ECM (1:4 hydration state) | 2D $^1$H-$^{13}$C INEPT-HETCOR | 256 | 3 | 83 | 108x185.15 | N/A |
| 2B, 3D (pink), 4A (pink), 4E, 6B (pink), S2D | HMW-HA (1:5, 1:8 hydration state) | 2D $^{13}$C-$^{13}$C TOBSY | 128 | 3 | 50 | 274x50.97 | 6 |
| 3D (green), 4A (green), S3, S4 | HMW-HA (1:0.5, 1:1 hydration state) | 2D $^{13}$C-$^{13}$C DARR | 16 | 3 | 83 | 732x24.53 | 8, 80 |

| Figure | Sample | Experiment | NS | RD (s) | TPPM (kHz) | $t_1$ evol. (□s) | Mixing (ms) |
|---|---|---|---|---|---|---|---|
| 2D | HMW-HA (1:5) | 2D $^{13}C$-$^{13}C$ INADEQUATE | 64 | 3 | 50 | 202x14 | N/A |
| S9 (top panel), S10A | HMW-HA (1:0.5, 1:1, 1;2, 1:5 hydration state) | $^{13}C$ DE $T_1$ relaxation | 1500 | 3.5 | 50 | N/A | N/A |
| S8 (top panel), S10B | HMW-HA (1:0.5, 1:1, 1;2, 1:5 hydration state) | $^{13}C$ DE $T_2$ relaxation | 2048 | 3.5 | 50* | N/A | N/A |
| S9 (bottom panel), S10C, | HMW-HA-ECM (1:0.5, 1:1, 1;2, 1:5 hydration state) | $^{13}C$ DE $T_1$ relaxation | 1500 | 3.5 | 50 | N/A | N/A |
| S8 (bottom panel), S10D | HMW-HA-ECM (1:0.5, 1:1, 1;2, 1:5 hydration state) | $^{13}C$ DE $T_2$ relaxation | 2048 | 3.5 | 50* | N/A | N/A |

*TPPM decoupling during echo time was 25kHz.

**Table S2: Chemical shift assignment of HMW-HA:** $^{1}H$ and $^{13}C$ chemical shift values of low (CP-based experiments) and high hydration state of HMW-HA (INEPT-based experiments).

| Carbon atom | $^{13}C$ chemical shift CP (1:0.5) (ppm) | $^{13}C$ chemical shift INEPT (1:5) (ppm) | $^{1}H$ chemical shift INEPT (1:5) (ppm) |
|---|---|---|---|
| **GlcNAc** | | | |
| C1a | 103.4 | 103.4 | 4.5 |
| C1b | - | 98.7 | - |
| C1c | 104.6 | - | - |
| C2a | 57.0 | 57.1 | 3.8 |
| C2b | 55.8 | 55.0 | - |
| C3a | 85.5 | 84.9 | 3.7 |
| C3b | 87.2 | - | - |
| C4a | 70.9 | 71.2 | 3.5 |
| C4b | 71.2 | 72.2 | 3.2 |
| C4c | - | 71.9 | 4.0 |
| C4d | - | 71.3 | 3.7 |
| C5a | 78.3 | 78.1 | 3.5 |
| C5b | 78.5 | 77.3 | - |
| C5c | - | 73.7 | 3.7 |
| C6a | 63.5 | 63.3 | 3.7 |
| C6b | 64.6 | 65.2 | 3.6 |
| C6c | - | 66.6 | 3.7 |
| C6d | - | 66.8 | 4.0 |
| C7a | 177.9 | - | - |
| C8a | 25.3 | 25.3 | 2.0 |
| **GlcA** | | | |
| C1’a | 106.8 | 105.8 | 4.5 |
| C1’b | 107.2 | 98.2 | - |
| C1’c | 107.2 | | |
| C2’a | 74.4 | 75.2 | 3.3 |
| C2’b | 75.7 | 75.3 | 3.6 |
| C3’a | 76.3 | 76.2 | 3.6 |
| C3’b | 75.7 | 75.2 | 3.6 |
| C4’a | 81.9 | 82.7 | 3.7 |
| C4’b | 87.7 | - | - |
| C5’a | 79.0 | 79.0 | 3.7 |
| C6’a | 177.6 | - | - |

**Table S3: Chemical shift assignment of HMW-HA-ECM:** $^{1}H$ and $^{13}C$ chemical shift values of HMW-HA-ECM in low (CP-based experiments) and high hydration state (INEPT-based experiments).

| Carbon atom | $^{13}C$ CP chemical shift (1:1.5) (ppm) | $^{13}C$ chemical shift INEPT (1:4) (ppm) | $^{1}H$ chemical shift INEPT (1:4) (ppm) |
|---|---|---|---|
| **GlcNAc** | | | |
| C1a | 103.4 | 103.5 | 4.5 |
| C1b | - | 98.7 | 4.6 |
| C1c | 104.6 | - | - |
| C1d | - | 101.0 | - |
| C2a | 57.0 | 57.1 | 3.8 |
| C2b | 55.8 | 55.0 | - |
| C2c | - | 56.7 | 4.2 |
| C3a | 85.5 | 84.9 | 3.7 |
| C3b | 87.2 | - | - |
| C4a | 70.9 | 70.8 | 3.5 |
| C4b | 71.2 | 72.2 | 3.9 |
| C4c | - | 71.9 | 4.0 |
| C4d | - | 71.3 | 3.7 |
| C4e | - | 72.5 | 3.4 |
| C5a | 78.3 | 78.0 | 3.5 |
| C5b | 78.5 | 77.3 | 3.2 |
| C5c | - | 73.9 | 3.8 |
| C6a | 63.5 | 63.3 | 3.7 |
| C6b | 64.6 | 65.2 | 3.6 |
| C6c | - | 66.1 | - |
| C6d | - | 66.8 | 4.0 |
| C6e | - | 64.9 | - |
| C7a | 177.9 | - | - |
| C8a | 25.3 | 25.3 | 2.0 |
| **GlcA** | | | |
| C1’a | 106.8 | 105.8 | 4.5 |
| C1’b | 107.2 | 98.2 | - |
| C1’c | 107.2 | | |
| C2’a | 74.4 | 75.2 | 3.3 |
| C2’b | 75.7 | 75.3 | 3.6 |
| C3’a | 76.3 | 76.2 | 3.6 |
| C3’b | 75.7 | 75.7 | 3.6 |
| C4’a | 81.9 | 82.7 | 3.7 |
| C4’b | 87.7 | - | - |
| C5’a | 79.0 | 79.0 | 3.7 |
| C6’a | 177.6 | - | - |

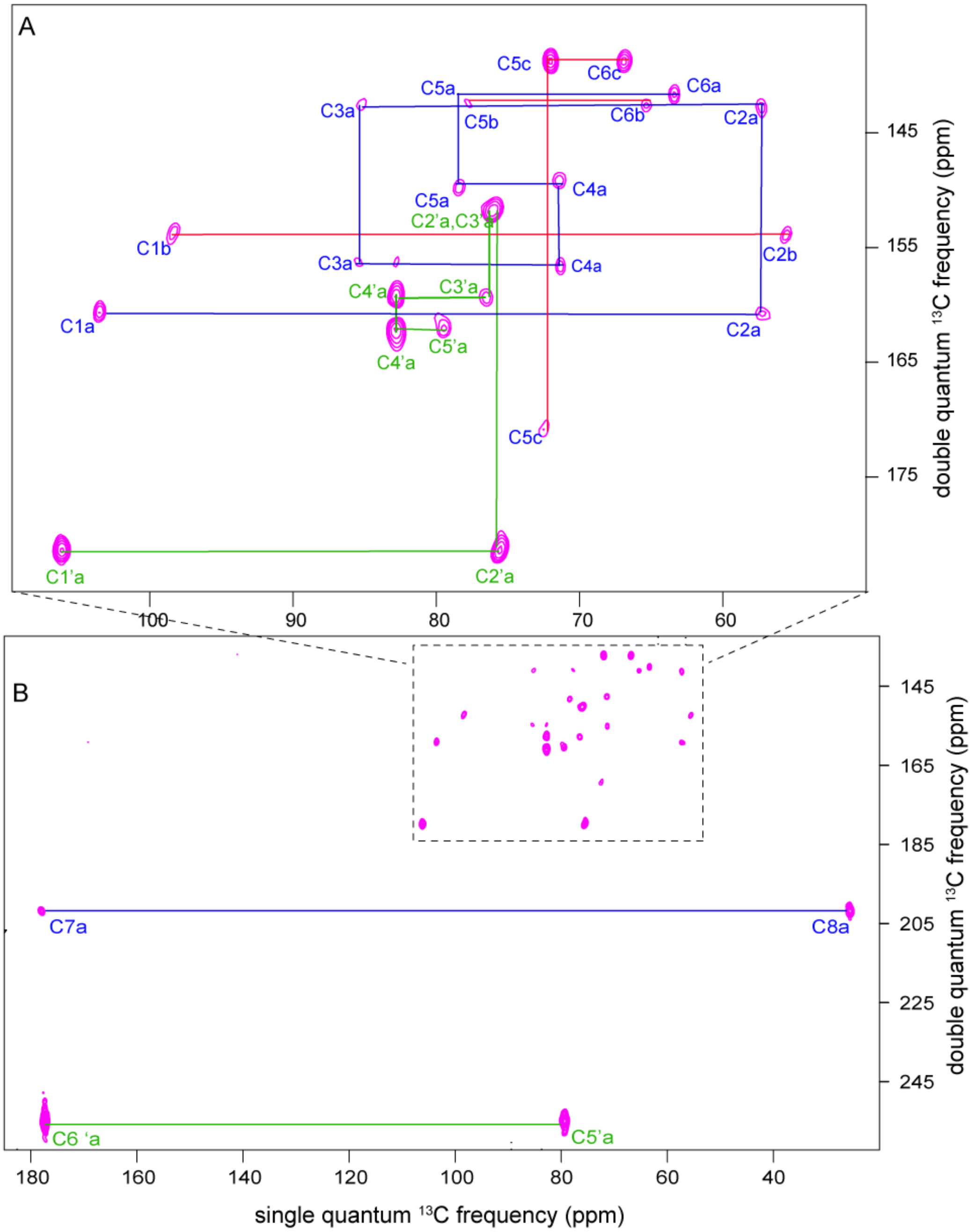


**Figure S1. 2D $^{13}C$ MAS NMR of $^{13}C$-HMW-HA via $^{13}C$ DQ-SQ INADEQUATE spectroscopy.** (A) Zoomed region from the 2D $^{13}C$ INADEQUATE spectrum of HA:$D_2O$ 1:5 (w/v), showing the ring, side chain and anomeric carbons of both moieties. Red lines indicate extra conformations of GlcNAc moiety. (B) The carbonyl and methyl peaks are visible in the full range spectrum, marked for GlcA (green line) and GlcNAc (blue line).

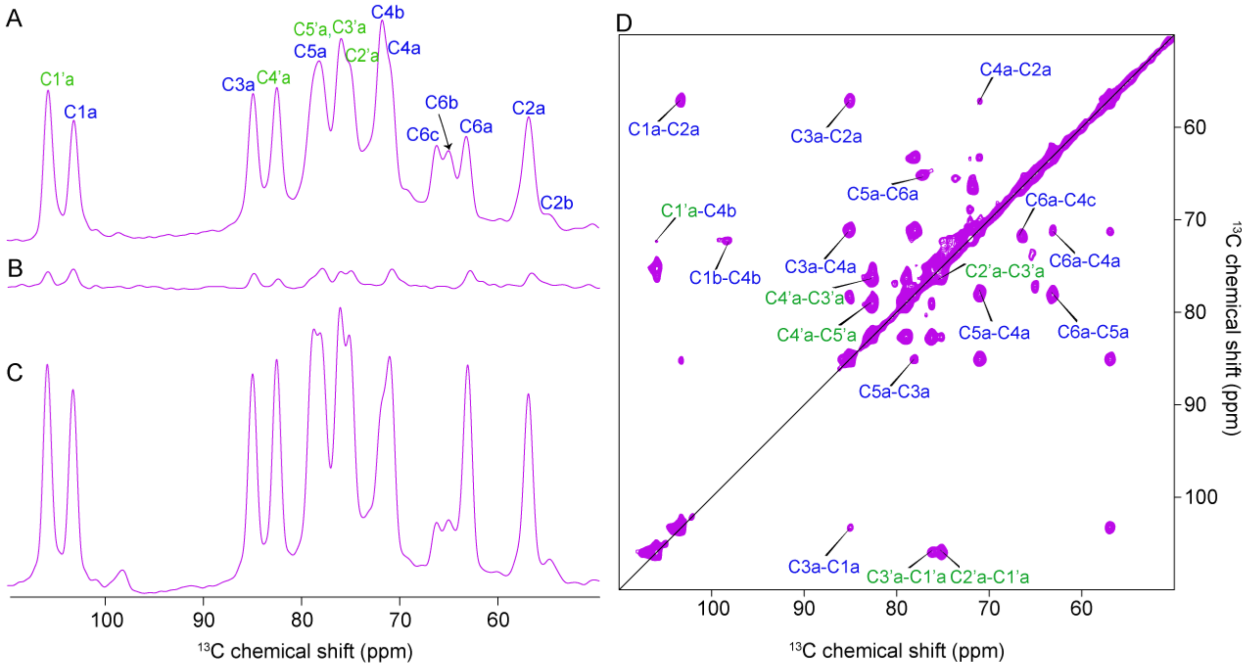


**Figure S2. 1D and 2D $^{13}$C-$^{13}$C INEPT-TOBSY spectrum of HMW-HA (1:8 hydration ratio, w/v).** (A) 1D INEPT (B) CP and, (C) DE spectra. (D) 2D $^{13}$C-$^{13}$C INEPT-TOBSY spectrum Measured with 6ms TOBSY mixing time, 10kHz MAS and temperature set to 277K. Green labels represent the carbons from the GlcA and blue ones are from the GlcNAc moiety.

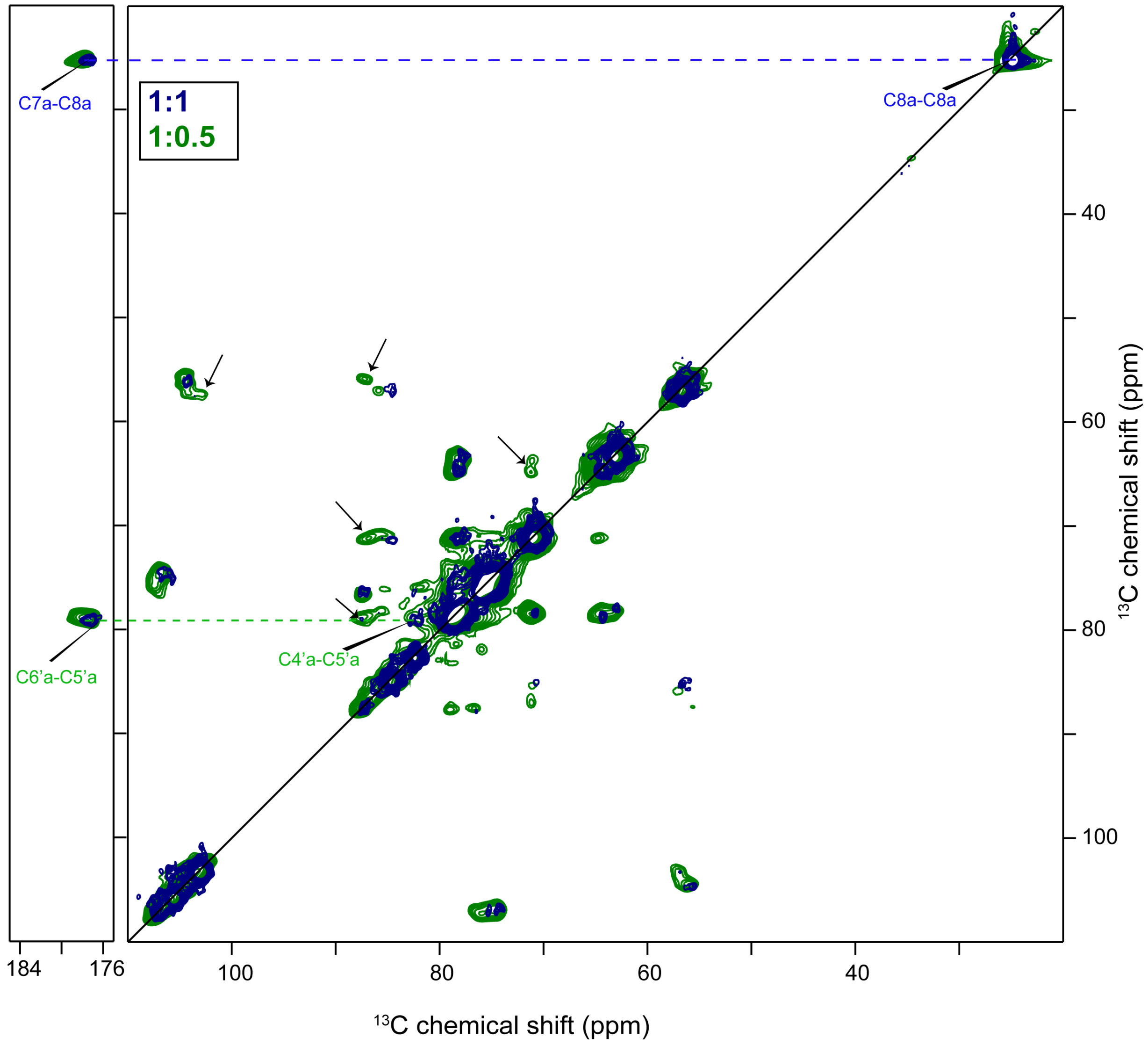


**Figure S3. Overlayed 2D $^{3}$C-$^{13}$C CP-based DARR ssNMR spectra, comparing HMW-HA 1:1 (w/v) with HMW-HA 1:0.5 hydration ratio (w/v).** The spectra reveal a difference in the number of observed extra conformations in HA as a function of its hydration level. Increases in hydration level (blue spectrum) result in enhanced flexibility, such that carbons which are more flexible become invisible in this CP-based NMR spectrum (indicated with arrows). The methyl (C8), carbonyl (C7) and carboxyl regions (C6') are also labeled. Dashed lines indicate connectivities with the carbonyl/carboxyl and methyl regions.

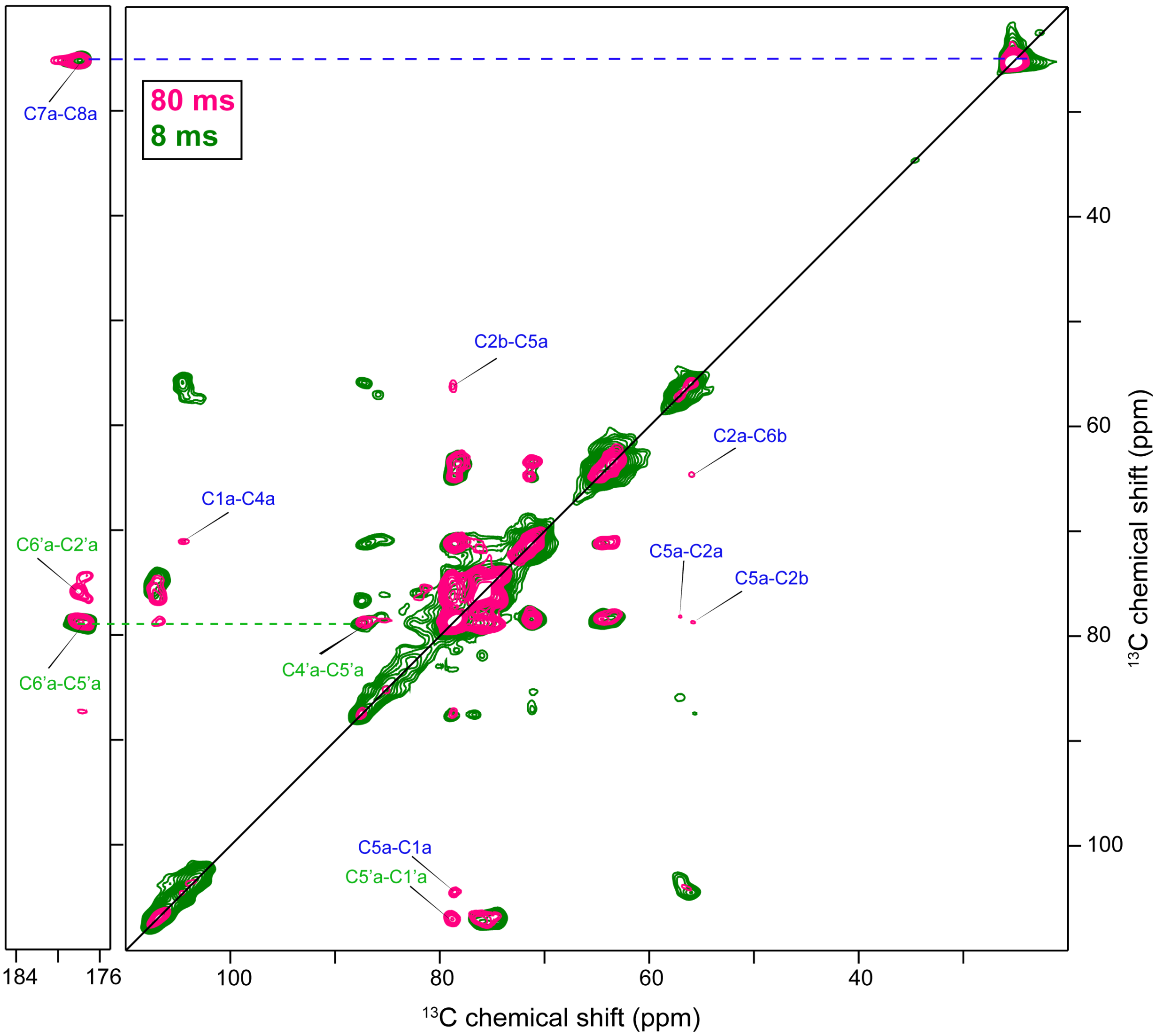


**Figure S4. Overlay of 2D $^{13}C$-$^{13}C$ DARR spectra showing short- and long-range interactions within $^{13}C$ HMW-HA in the low hydration state (1:0.5, w/v).** The overlayed spectra reflect short (8ms) and long (80ms) DARR $^{13}C$-$^{13}C$ polarization transfer times, as indicated. Long-range interaction peaks are labeled with text labels that are color coded for the two monosaccharides in HA (GlcA in green and GlcNAc in blue). The carbonyl region is also shown at the left. Dashed lines indicate one bond $^{13}C$ connectivities between carbonyl and methyl, as well as carboxyl and ring carbon.

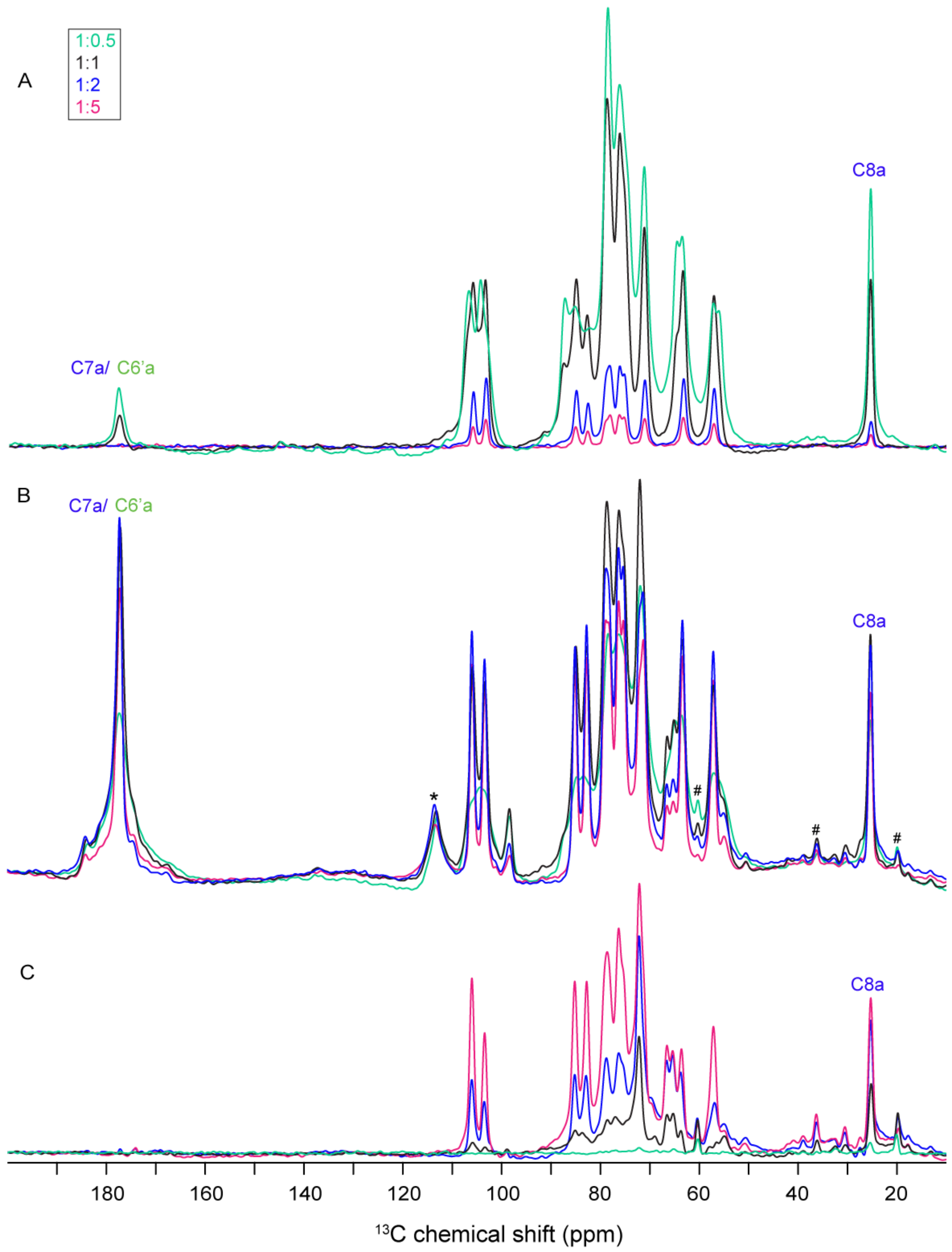


**Figure S5. Full 1D $^{13}$C ssNMR spectra of HMW-HA**. (A) 1D $^{13}$C CP, (B) 1D $^{13}$C DE and (C) 1D $^{13}$C INEPT. The methyl, carbonyl, and carboxyl carbons are indicated as C8a, C7a and C6'a, respectively. In DE spectra (panel B), peaks labeled with an asterisk (*) are from the sample spacer, while hash (#) labels indicate impurities in the sample.

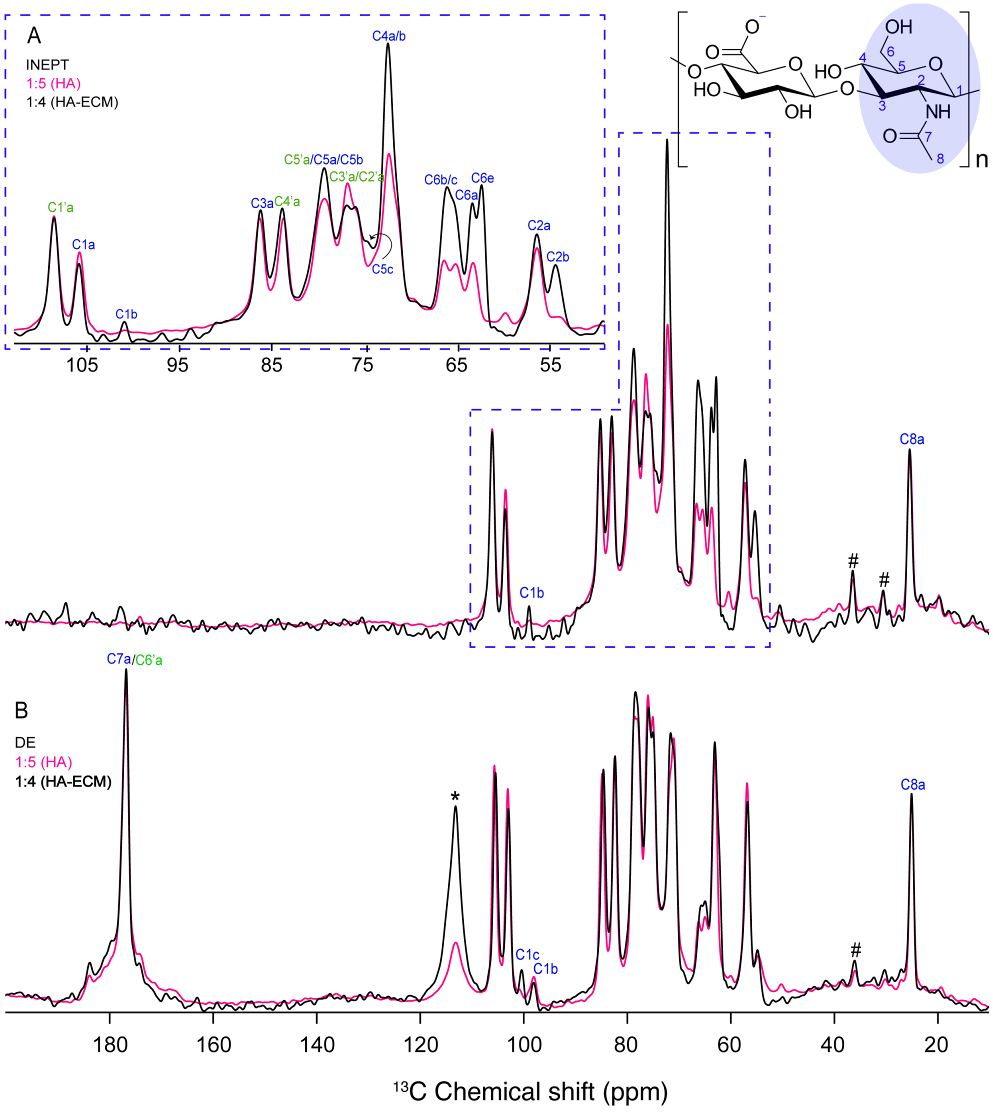


**Figure S6. Overlayed 1D $^{13}$C INEPT and DE ssNMR spectra of high hydration state of HMW-HA (1:5, w/v) and HMW-HA-ECM (1:4, w/v).** (A) Overlayed INEPT spectra of HMW-HA (pink) and HMW-HA-ECM (black). Dashed box in panel A is enlarged in the top left inset, highlighting spectral differences between HMW-HA and HA-ECM complex. In the HMW-HA-ECM complex, most of the ring and side chain carbons from the GlcNAc moiety show the enhanced intensity and the appearance of extra form of the anomeric carbon (C1b), indicating the increased flexibility compared to the GlcA moiety. (B) Overlayed DE spectra from the same samples. The shaded blue circle (top right) indicates the GlcNAc moiety in the chemical structure of HMW-HA, which is characterized by higher flexibility in HA chain in the HMW-HA-ECM complex. Blue and green labels represent carbons from GlcNAc and GlcA. Peaks labelled with hash sign are from the impurities and the peak indicated with an asterisk is from the sample spacer.

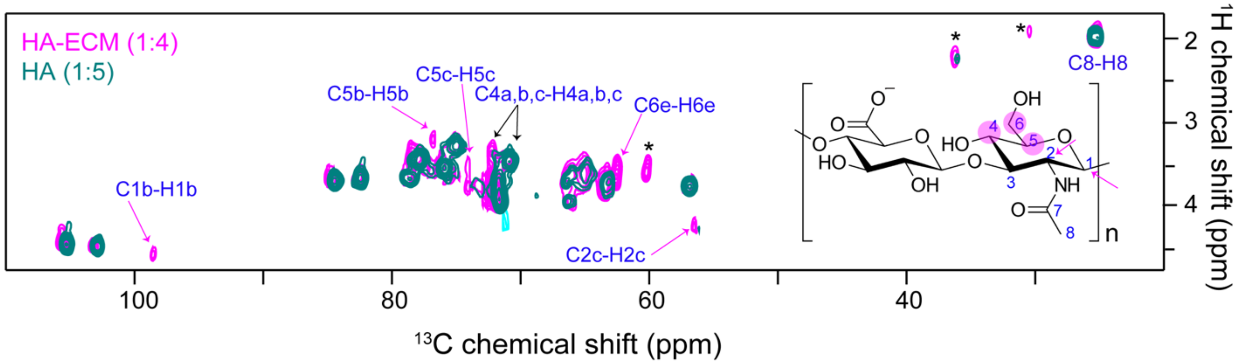


**Figure S7. Overlayed 2D $^1$H, $^{13}$C HETCOR spectra of highly hydrated state of both HMW-HA (1:5, w/v) and HMW-HA-ECM (1:4, w/v)**. Additional peaks, denoted by arrows, emerge in the HMW-HA-ECM sample. Specifically, carbons C5 and C6 exhibit extra peaks (pink arrows), while carbons C1 and C4 demonstrate chemical shift perturbations, marked by black arrows, in the HMW-HA-ECM spectrum. The most significantly impacted carbons are emphasized with solid (pink) circles and arrows on the chemical structure of HMW-HA (on the right). Peaks labeled with asterisks originate from ECM components.

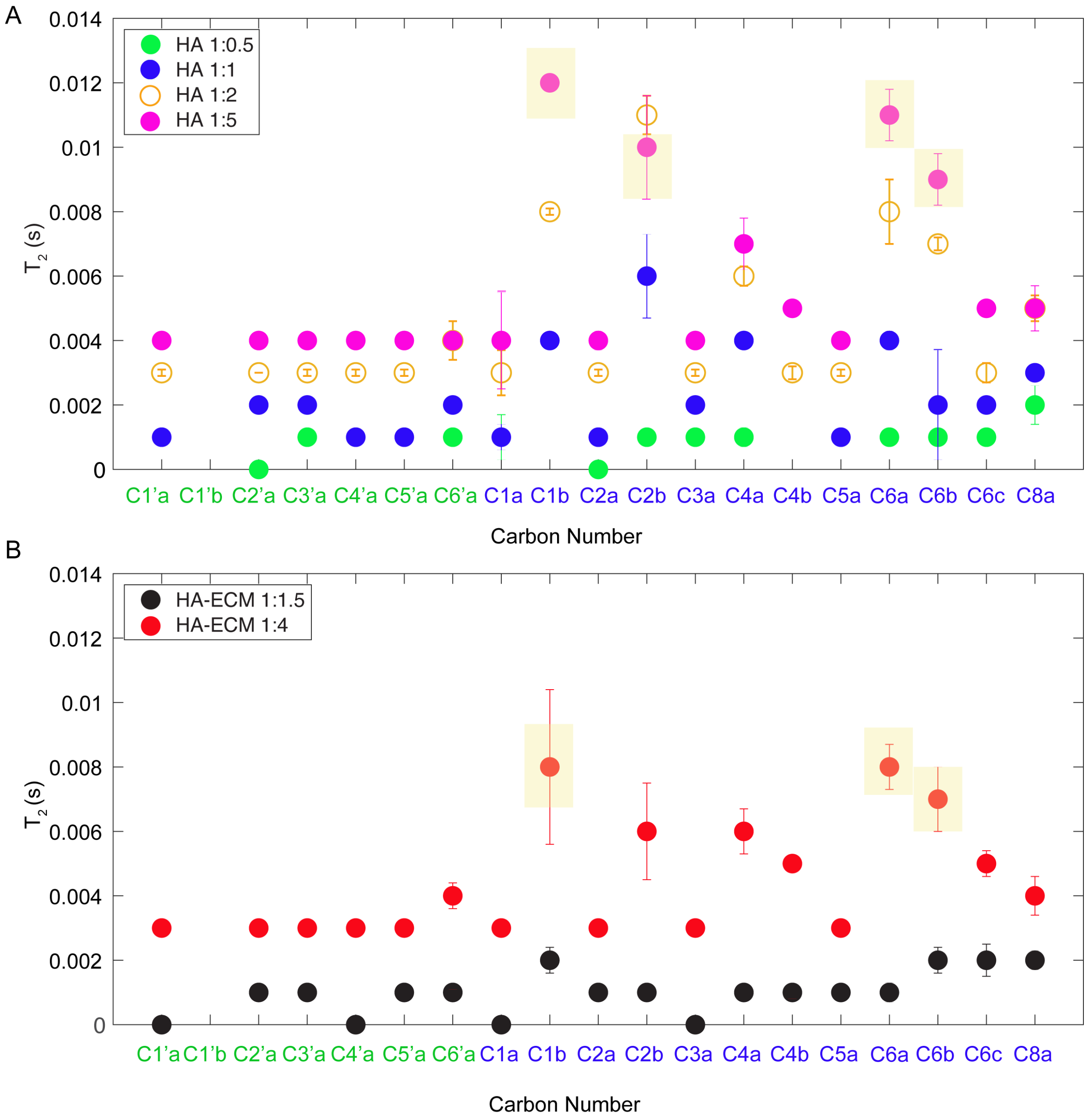


**Figure S8. $T_2$ relaxation plots of HMW-HA (A) and HMW-HA-ECM (B) showing hydration dependent global and local dynamic changes.** Increased hydration levels show higher $T_2$ values indicating enhanced dynamics of HA chain. At most conditions the $T_2$ values are similar for most of the HA chain carbons, indicating that a similar flexibility throughout the molecule. Yellow boxes highlight a subset of carbons which show higher $T_2$ values, which identifies locally increased flexibility affecting those atoms. On the horizonal axis, green labels represent carbon numbers from the GlcA moiety while blue labels represent carbons from the GlcNAc moiety.

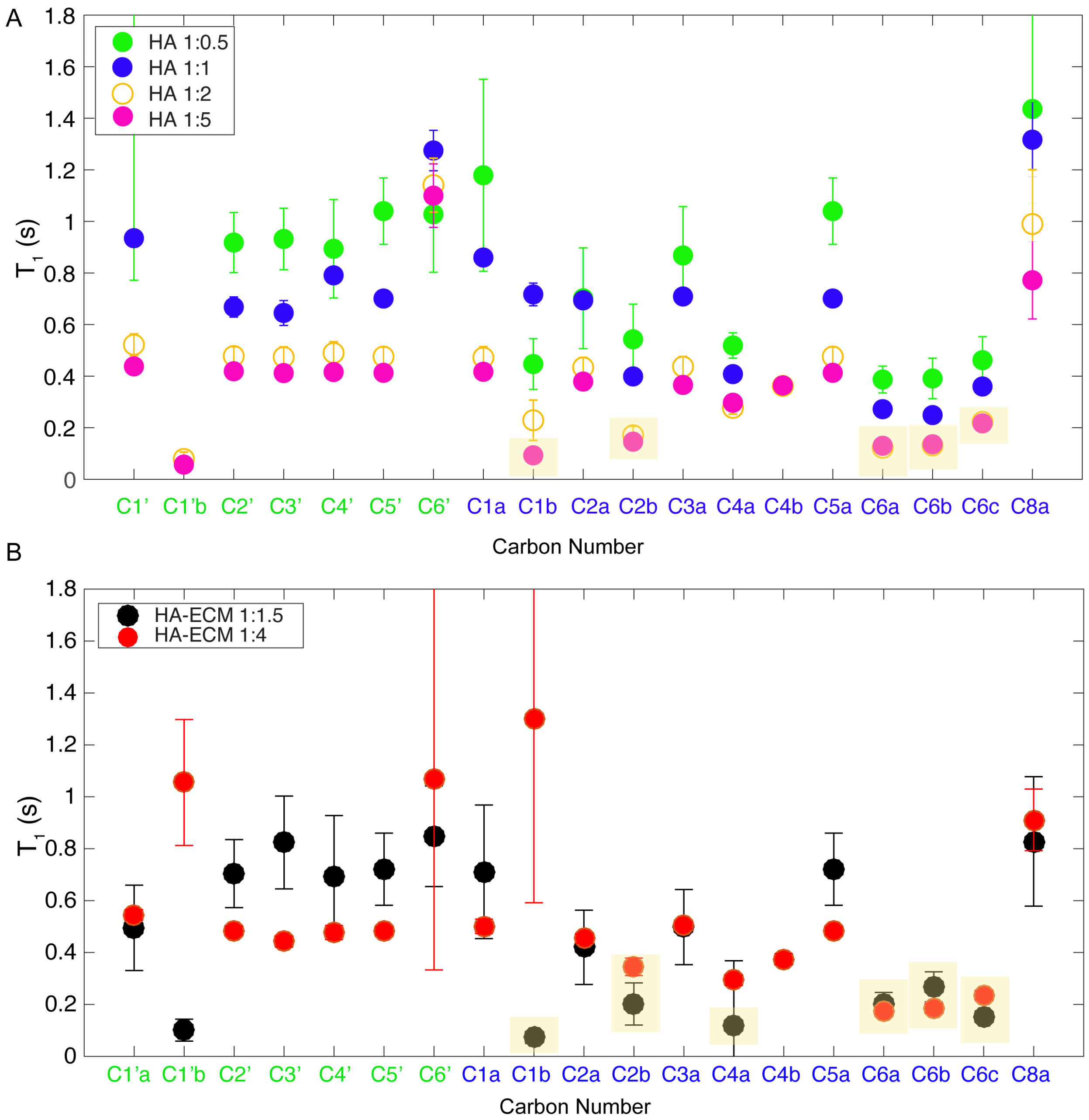


**Figure S9. $T_1$ relaxation plots of HMW-HA (A) and HMW-HA-ECM (B) showing hydration dependent global and local dynamic changes.** Increased hydration levels show lower $T_1$ values indicating enhanced flexibility of HA chain. Most carbons (at a given condition) have similar T1 values, indicating that a similar mobility of the entire macromolecule. However, selected carbon peaks show lower $T_1$ values compared to the rest of the chain, as marked with yellow boxes. On the horizonal axis, the green labels represent carbon numbers from the GlcA moiety while blue labels represent carbons from the GlcNAc moiety.

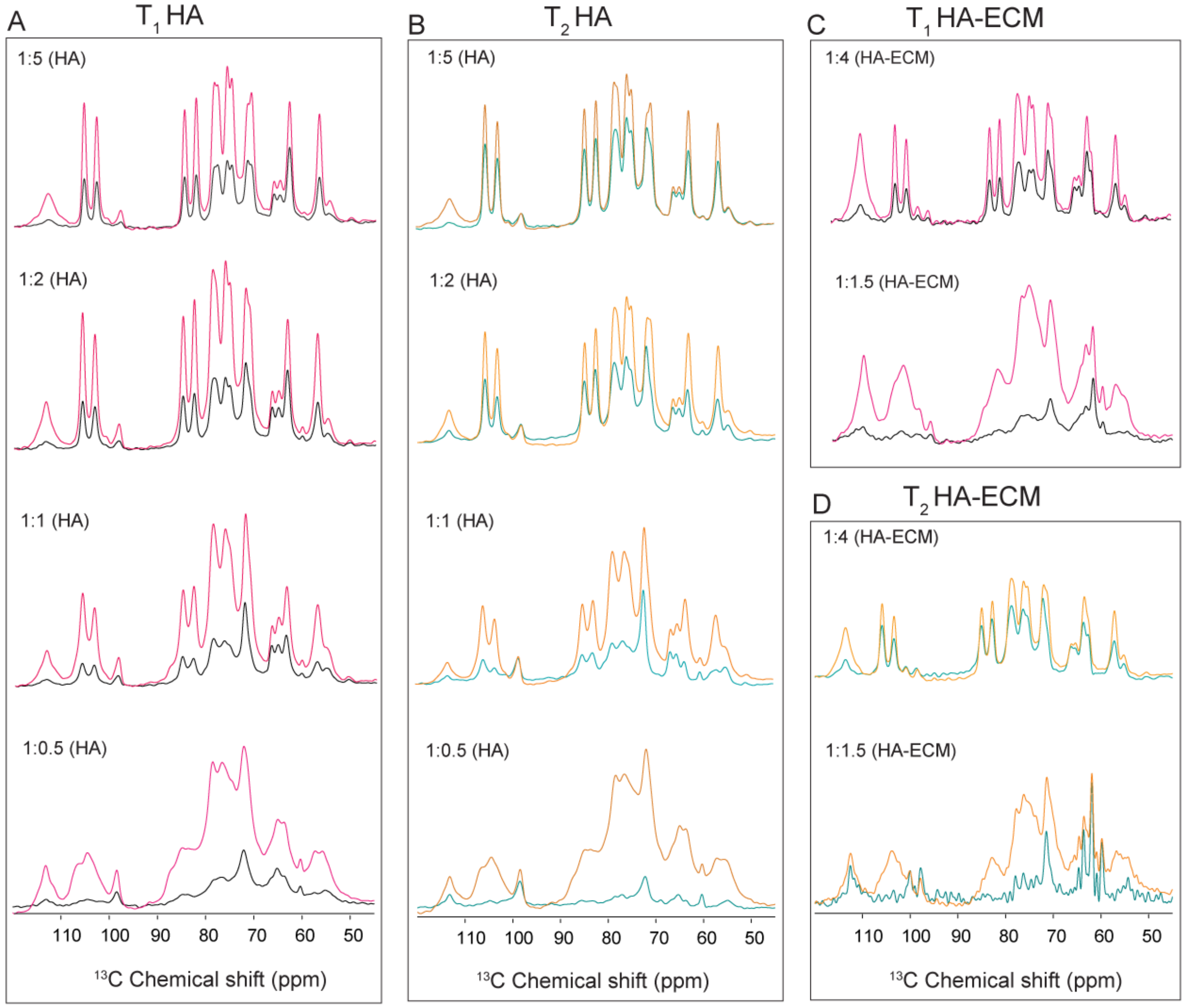


**Figure S10. Representative $^{13}C$ $T_1$ and $T_2$ ssNMR spectra.** (A) Overlayed $T_1$ relaxation spectra of various hydration levels of HMW-HA with two different recovery delays, 250ms (black) and 10s (pink). (B) Overlayed $T_2$ relaxation spectra of various hydration levels of HMW-HA with two different Hahn echo times, 0ms (orange) and 16ms (teal color). (C, D) Analogous $T_1$ and $T_2$ relaxation measurement spectra for $^{13}C$-HMW-HA complexed with unlabeled ECM components. The employed pulse sequence is based on an initial generation of $^{13}C$ signal by direct excitation.

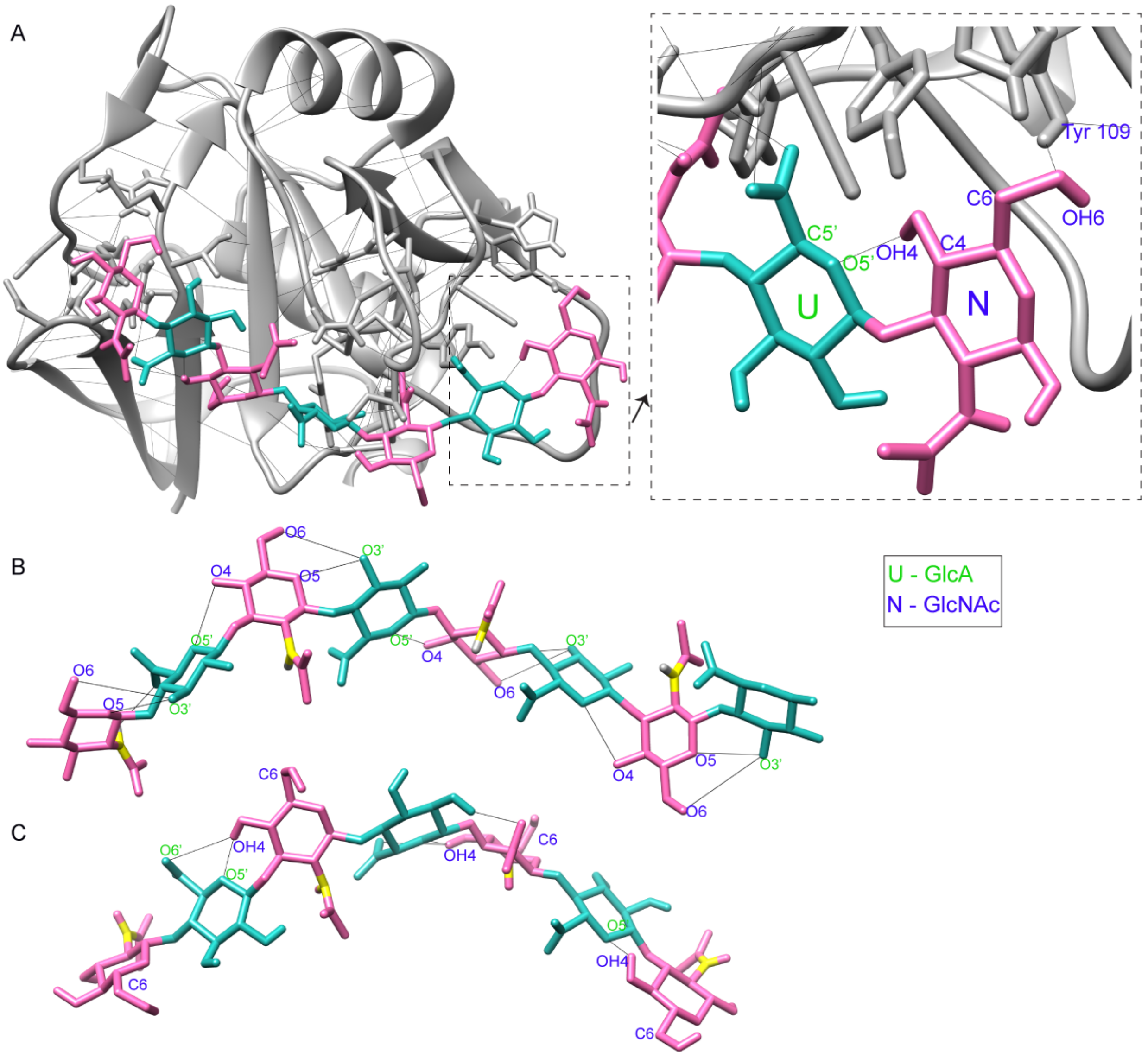


**Figure S11. Atomic interactions of HA octamer with HA-binding protein.** (A) Ribbon diagram of the HA binding domain of mouse CD44 bound to HA octasaccharide (PDB-ID 2JCQ) [1]. Dashed box shows zoomed-in region of protein residues binding to the HA oligomer. (B) HA conformation from a combination of MD simulation and solution NMR (PDB-ID, 2BVK) [2]. (C) Conformation of HA when bound to CD44 HABD (same as panel A, the protein structure was hidden to visualize the HA conformation in the bound form). In panel B and C, solid lines show hydrogen bonds formed between oxygen atoms which are attached to carbon C6, C4, C5 in GlcNAc, and C3', C5' in GlcA moieties.

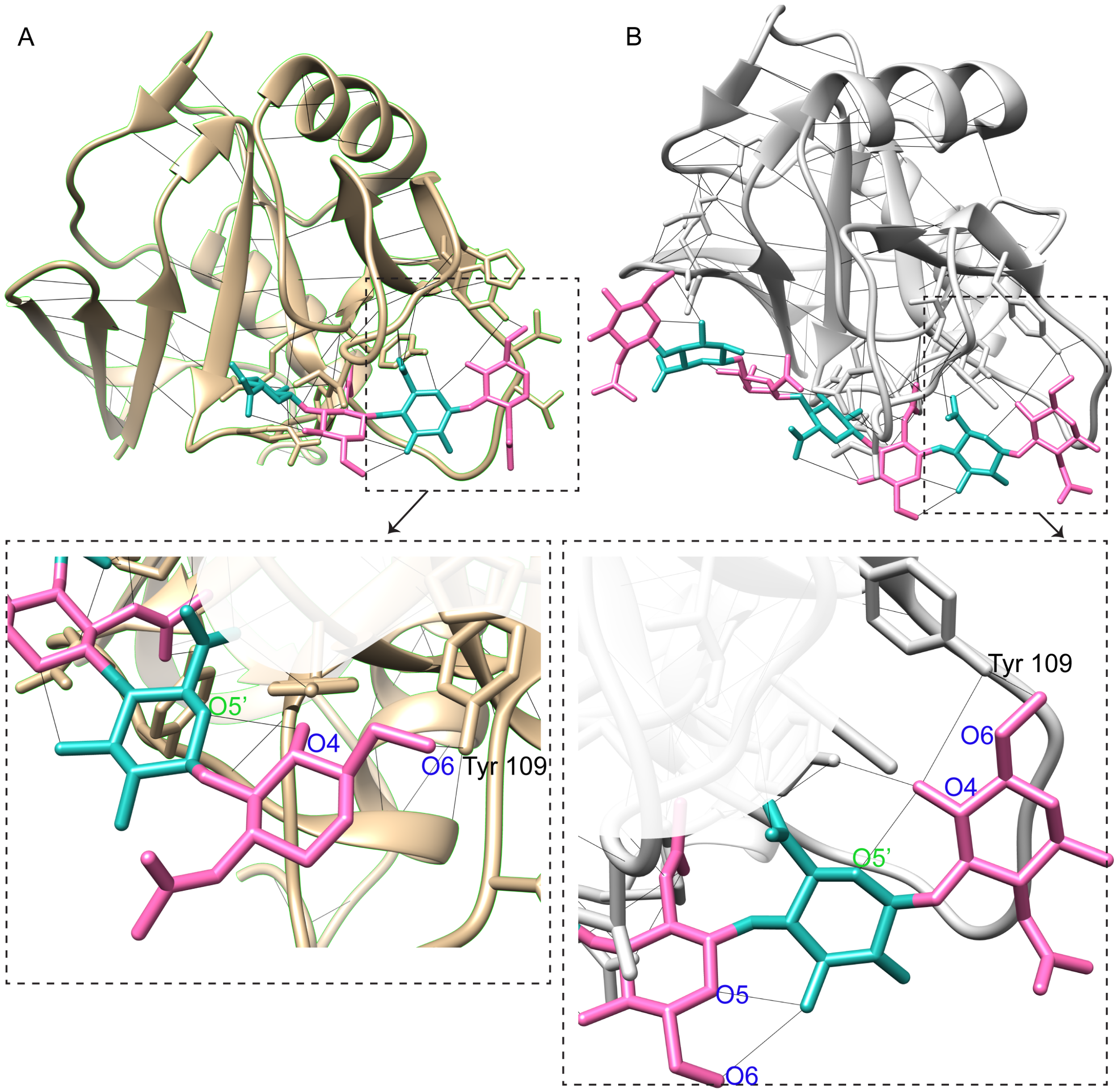


**Figure S12. Molecular perspectives of HA interactions in known structures of HA-binding proteins.** (A) Ribbon diagram of human CD44 HABD (gray) with bound HA tetrasaccharide (cyan, pink, PDB-ID 4MRD) [3], and of (B) mouse CD44 (type B complex) with bound HA octasaccharide (cyan and pink, PDB-ID 2JCR) showing atomic interactions of HA [4]. The carbons 6 and 4 are labeled to show interactions with the proteins. In both cases bound HA-protein structure shows the formation of a hydrogen between the O6 from the GlcNAc moiety with the hydroxyl oxygen of Tyr 109. In dashed boxes the zoomed in regions of HA-protein bound region are shown.
